# Nucleotide imbalance decouples cell growth from cell proliferation

**DOI:** 10.1101/2021.12.06.471399

**Authors:** Frances F. Diehl, Teemu P. Miettinen, Ryan Elbashir, Christopher S. Nabel, Scott R. Manalis, Caroline A. Lewis, Matthew G. Vander Heiden

## Abstract

Nucleotide metabolism supports RNA synthesis and DNA replication to enable cell growth and division. Nucleotide depletion can accordingly inhibit cell growth and proliferation, but how cells sense and respond to changes in the relative levels of individual nucleotides is unclear. Moreover, the nucleotide requirement for biomass production changes over the course of the cell cycle, and how cells coordinate differential nucleotide demands with cell cycle progression is also not well understood. Here we find that excess levels of individual nucleotides can inhibit proliferation by disrupting the relative levels of nucleotide bases needed for DNA replication. The resulting purine and pyrimidine imbalances are not sensed by canonical growth regulatory pathways, causing aberrant biomass production and excessive cell growth despite inhibited proliferation. Instead, cells rely on replication stress signaling to survive during, and recover from, nucleotide imbalance during S phase. In fact, replication stress signaling is activated during unperturbed S phases and promotes nucleotide availability to support DNA replication. Together, these data reveal that imbalanced nucleotide levels are not detected until S phase, rendering cells reliant on replication stress signaling to cope with this metabolic problem, and disrupting the coordination of cell growth and division.

## Introduction

Most proliferating cells double each component of their mass over the course of the cell cycle, leading to metabolic demands that shift to enable biosynthetic processes specific to different cell cycle phases^1, 2^. A notable example is the requirement for nucleotides, which is particularly high in proliferating cells. Cells must acquire sufficient levels of each nucleotide species both for RNA synthesis and biomass production, and to ensure efficient and accurate DNA replication during S phase.

The enzyme ribonucleotide reductase (RNR) is responsible for dNTP production, converting ribonucleoside diphosphates to deoxyribonucleoside diphosphates. RNR inhibition impairs DNA replication and induces a replication stress response^3^, indicating that a substrate-level limitation for dNTPs can impede DNA synthesis. In *S. cerevisiae*, insufficient dNTPs at the onset of S phase can activate a replication stress response in unperturbed cells^4^, suggesting that endogenous dNTP levels are within a range that can become limiting for eukaryotic cells. Moreover, mutations in RNR that lead to depletion of specific dNTP species can also slow S phase progression in budding yeast^5^, underscoring the importance of maintaining appropriate levels of dNTPs to support DNA replication for cell division.

Proliferating cells also require nucleotides for the synthesis of rRNA and mRNA needed for biomass production. RNA production contributes to biomass both directly, as RNA comprises the vast majority of nucleic acid in cells^6^, and indirectly, as it is essential for transcription and translation of proteins. This underscores the importance of nucleotide acquisition not only for cell cycle progression and division, but also for the production of new biomass to support cell growth. Moreover, this raises the question of how cells coordinate differential needs for nucleotide species in supporting cell growth and enabling genome replication specifically during S phase.

Cells have evolved signaling networks that match growth with metabolic capacity and are conserved across species, enabling metabolism to support cell proliferation across a range of environmental conditions. Growth control pathways coordinate responses to stress conditions, as well as the availability of nutrients needed for cell proliferation^1^. Cellular responses to nutrient limitation generally arrest growth and downregulate macromolecule biosynthesis to preserve cellular resources until conditions improve to enable proliferation^1, 7, 8^. Growth signaling networks play a role in controlling nucleotide metabolism, both by coordinating the production and breakdown of RNA, and by regulating *de novo* nucleotide synthesis. For example, the mTORC1 substrate p70 S6 kinase directly phosphorylates and stimulates a key enzyme in pyrimidine synthesis^9^, and mTORC1 signaling also enhances production of one-carbon substrates for purine synthesis^10^. Nucleotide availability can also be an important input for growth control pathways, and mTORC1 activity is regulated by purine levels^11, 12^. In cells with defective autophagy, providing nucleotides alone allows survival in starvation conditions^13^, highlighting the importance of maintaining nucleotide homeostasis for cellular fitness.

Each nucleotide species has distinct roles in cell metabolism. In addition, levels of each nucleotide vary over a wide range of intracellular concentrations, and will be differentially affected by environmental conditions^6^. Extracellular nucleotide availability can vary based on physiological context^1, 14^, and while many cells salvage available nucleobases and nucleosides rather than synthesize them *de novo*, the relative environmental scarcity of these species means that many cells rely on *de novo* synthesis to fulfill at least part of their nucleotide demands. Both purine and pyrimidine production involve multiple metabolic pathways, which can be differentially affected by nutrient limitations or redox perturbations^1, 15–17^. Thus, the availability of different environmental nutrients, including nucleotides, can affect relative levels of individual nucleotides in cells. However, it remains less clear how cells maintain balanced nucleotides for shifting demands throughout the cell cycle, and whether cells sense the relative availability of specific nucleotide species.

Here we show that purine and pyrimidine imbalances inhibit cell proliferation, but nucleotide imbalances are not sensed by canonical growth regulatory pathways. Rather, cells continue to grow despite nucleotide imbalance and enter S phase, leading to activation of replication stress signaling as a protective response to imbalanced nucleotides. Moreover, replication stress signaling promotes nucleotide availability during unperturbed S phases, suggesting that cells rely on sensing replication stress downstream of nucleotide levels to maintain nucleotide balance during normal proliferation.

## Results

### Individual nucleotide precursors can inhibit cell proliferation

Proliferating cells must adapt metabolism to produce biomass from available nutrients^2^. Thus, disrupting pro-growth signaling pathways, limiting amino acid availability, blocking mitochondrial respiration, and inhibiting nucleotide synthesis all can inhibit cell proliferation (Fig. 1a). Obtaining nucleotides for efficient RNA and DNA synthesis can be particularly limiting for cell proliferation^18–20^. Indeed, pharmacological inhibition of purine production with lometrexol (LTX) or pyrimidine production with brequinar (BRQ) depletes total purine or pyrimidine levels and blocks proliferation, consistent with previous studies^11, 21, 22^ (Fig. 1a and Extended Data Fig. 1a). Intriguingly, thymidine treatment has long been used to arrest and synchronize cells; however, the proximal mechanisms of how this arrest is achieved, and whether this has broader implications for the regulation of nucleotide homeostasis, is unclear. Moreover, thymidine exists uniquely in the deoxyribonucleotide pool, and it is unknown whether perturbations to ribonucleotide pools are equally detrimental. To investigate this, we provided cells with individual nucleobases and nucleosides, which can be salvaged to produce nucleotides. Nucleotide salvage preserves metabolic substrates that would otherwise be needed for *de novo* nucleotide synthesis. However, we found that in sufficient amounts, single nucleotide supplementation impaired proliferation (Fig. 1b-f). This effect was dose-titratable, with increasing concentrations of single nucleotides completely preventing cell proliferation.

**Figure 1.**
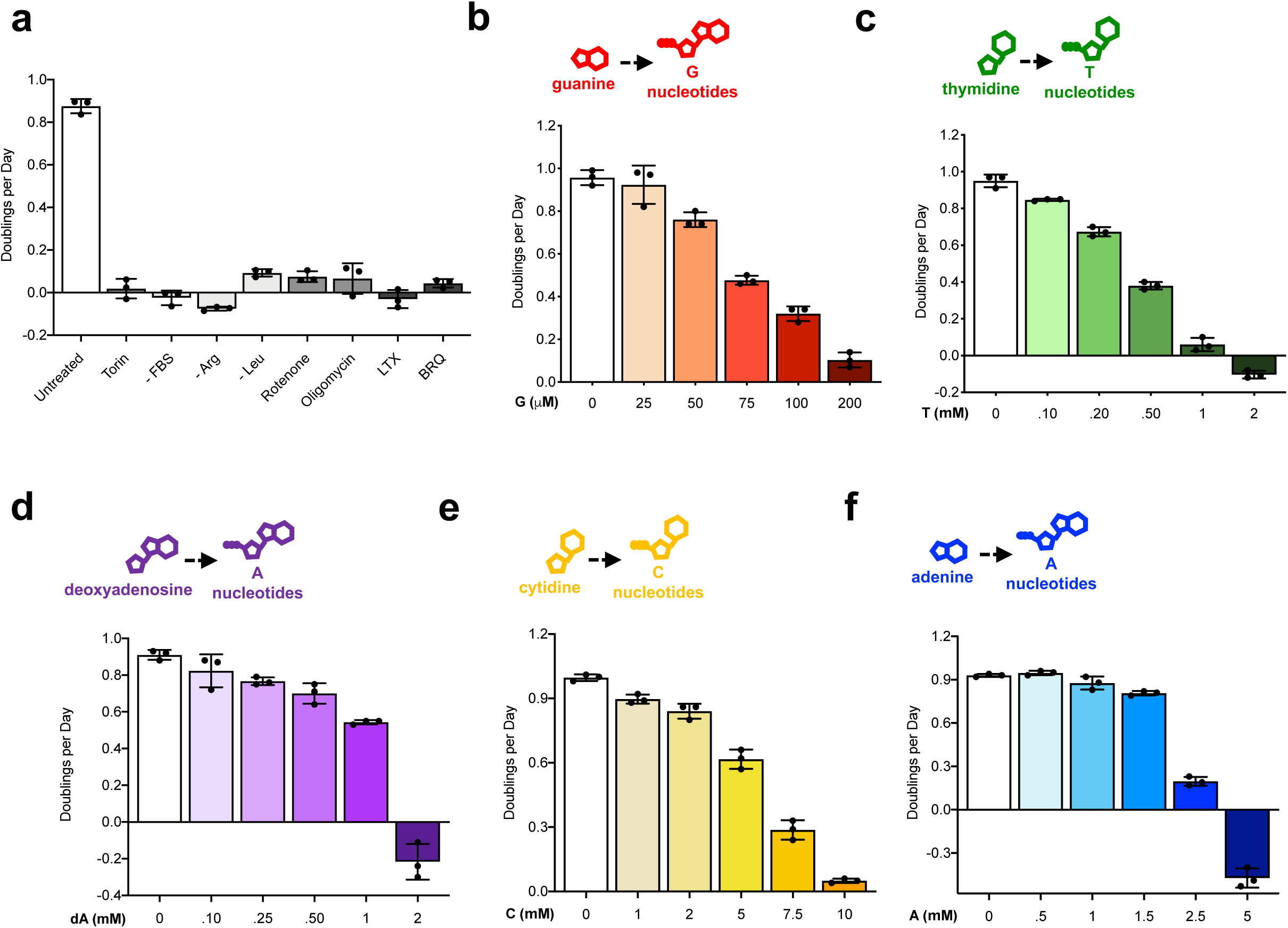
Excess individual nucleotides can impair proliferation. **a**, Proliferation rates of A549 cells cultured in standard conditions (Untreated) or with 1 µM Torin1, without serum (-FBS), without arginine (-Arg), without leucine (-Leu), with 100 nM rotenone, with 5 nM oligomycin, with 1 µM lometrexol (LTX), or with 1 µM brequinar (BRQ). **b**-**f**, Proliferation rates of A549 cells treated with the indicated concentration of guanine (G), thymidine (T), deoxyadenosine (dA), cytidine (C), or adenine (A). Each of these nucleobases/nucleosides can be salvaged to produce intracellular nucleotides as shown. Data are presented as mean +/- SD of 3 biological replicates.

Nucleotide species are maintained within different intracellular concentration ranges^23^. Expression of nucleotide salvage and synthesis enzymes, as well as transporters, also varies across cells and thus might be expected to affect sensitivity to individual nucleotide addition. Consistent with this, different cells had differential sensitivity to each nucleotide, although proliferation could be inhibited with precursor addition in all cells tested, including non-transformed cells (Extended Data Fig. 1b,c). Interestingly, deoxycytidine at concentrations up to 14 mM was the only precursor tested that did not block proliferation in these cells (Extended Data Fig. 1d). Because most cells exhibited greatest sensitivity to guanylate nucleotide precursors (Extended Data Fig. 1e), we focused further mechanistic studies on understanding the effects of guanine supplementation. Importantly, a functional salvage pathway was needed for the corresponding nucleotide precursor to inhibit proliferation: cells deficient for APRT and HPRT, the enzymes that salvage adenine and guanine, were not sensitive to these precursors, and thymidine kinase-deficient cells were unaffected by addition of thymidine (Extended Data Fig. 1b,f).

### Nucleotide salvage resulting in nucleotide imbalance impairs proliferation

We reasoned that salvage of single nucleobases/nucleosides might perturb the relative levels of nucleotide species, and tested this hypothesis by measuring intracellular nucleotide levels with and without guanine addition. Guanine supplementation increased intracellular pools of guanylate nucleotides (GTP/GDP/GMP). Unexpectedly, guanine also decreased intracellular levels of adenylate nucleotides (ATP/ADP/AMP) (Fig. 2a and Extended Data Fig. 2a). These data suggest that guanine salvage disrupts normal purine balance by increasing the ratio of guanylate (G) to adenylate (A) nucleotides. Notably, providing exogenous adenine together with guanine restored the balance of intracellular G and A nucleotides (Fig. 2a).

**Figure 2.**
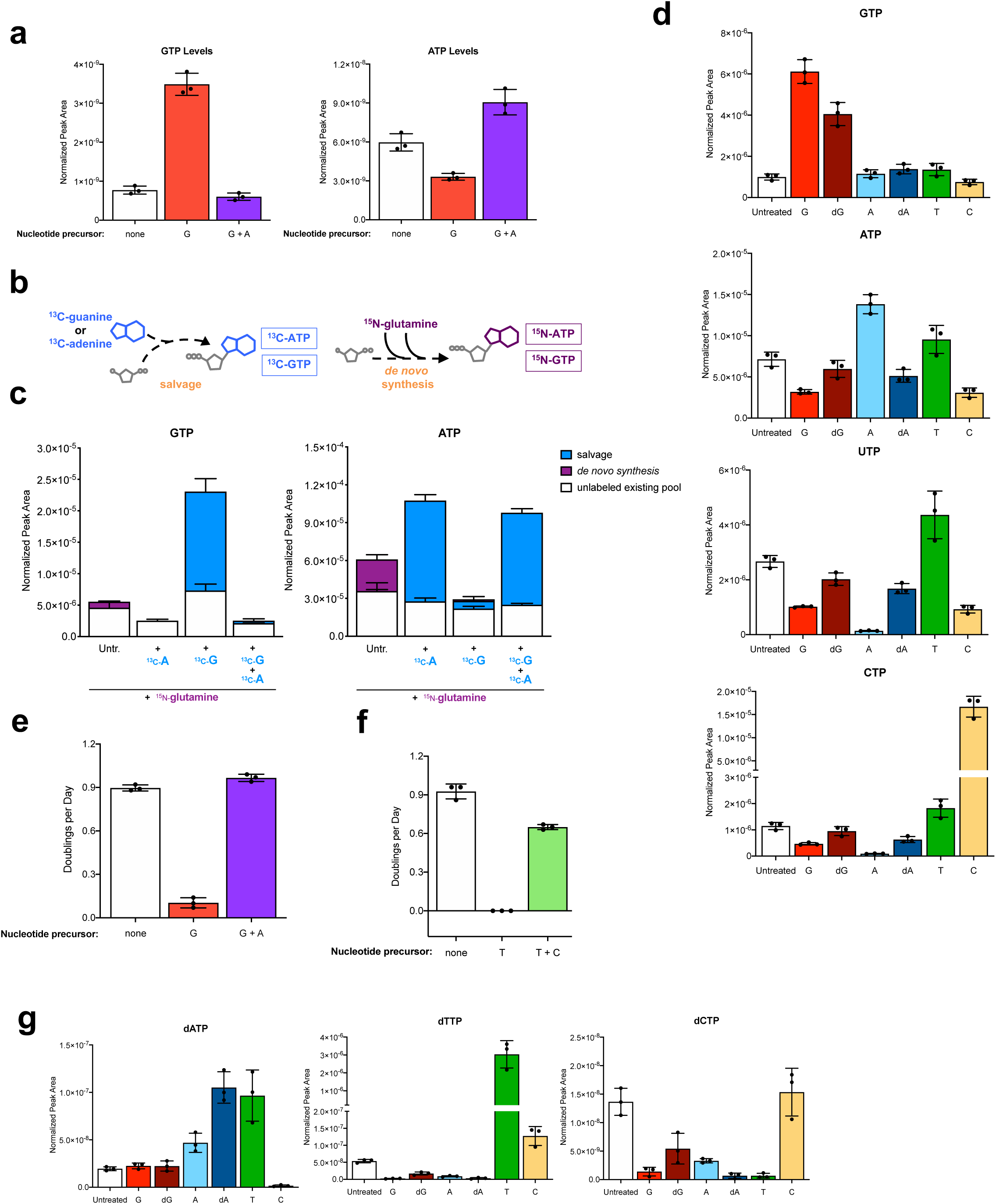
Nucleotide salvage leading to imbalanced nucleotide pools inhibits cell proliferation. **a**, GTP and ATP levels in A549 cells cultured in standard conditions (none) or treated for 24 hours with 200 µM guanine (G) with or without 200 µM adenine (A) as indicated. **b**, Schematic showing how stable isotope tracing was used to determine the source of intracellular purines. Salvage of ^13^C- guanine or ^13^C-adenine produces ^13^C-labeled GTP and ATP. The ^15^N label from amide-^15^N-glutamine is incorporated in *de novo* purine synthesis, producing ^15^N-labeled ATP and GTP. **c**, Total levels and labeling of GTP and ATP in A549 cells cultured for 24 hours in media containing amide-^15^N-glutamine with or without 200 µM ^13^C-guanine and/or ^13^C-adenine as indicated. **d**, Intracellular nucleotide levels in A549 cells cultured in standard conditions or treated with 200 µM guanine (G), 20 µM deoxyguanosine (dG), 2.5 mM adenine (A), 1.5 mM deoxyadenosine (dA), 1 mM thymidine (T), or 10 mM cytidine (C) as indicated. **e**, Proliferation rate of A549 cells cultured in standard conditions (none) or treated with 200 µM G with or without 200 µM A. **f**, Proliferation rate of A549 cells cultured in standard conditions (none) or treated with 1 mM T with or without 1 mM C. **g**, Intracellular levels of dNTPs in A549 cells in standard culture conditions (Untreated) or treated with 200 µM G, 20 µM dG, 2.5 mM A, 1.5 mM dA, 1 mM T, or 10 mM C as indicated. All nucleotide levels were measured using LCMS. Data are presented as mean +/- SD of 3 biological replicates.

To understand how providing guanine depletes intracellular A nucleotides, we traced the fate of stable isotope-labeled metabolites to determine the contribution of both salvage and *de novo* synthesis to cellular purines. To assess *de novo* synthesis, we measured incorporation of amide-^15^N- glutamine into purines: the amide nitrogen of glutamine is incorporated during AMP and GMP synthesis such that purines labeled with ^15^N have been made *de novo*. We also used ^13^C-guanine and ^13^C-adenine to measure how salvage of these nucleobases contributes to intracellular purines: ^13^C-labeled purines are derived from the addition of guanine or adenine to a ribose-phosphate backbone (Fig. 2b). As expected, a subset of purine nucleotides in untreated cells were labeled with ^15^N, reflecting their production via *de novo* synthesis (Fig. 2c and Extended Data Fig. 2b). Providing ^13^C-adenine increased levels of A nucleotides, the majority of which were ^13^C-labeled and therefore derived from adenine salvage. Similarly, salvage of ^13^C-guanine accounted for the increase in G nucleotide levels upon guanine supplementation. Moreover, providing either ^13^C-adenine or ^13^C- guanine eliminated the contribution of *de novo* synthesis to both A and G nucleotide pools. This likely reflects known allosteric feedback regulation of purine synthesis enzymes: A and G nucleotides can inhibit the activity of both ribose-5-phosphate pyrophosphokinase and glutamine phosphoribosyl pyrophosphate amidotransferase, which catalyze the initial steps of *de novo* purine synthesis^6, 24, 25^. Aberrantly high levels of G nucleotides derived from guanine salvage can therefore inhibit *de novo* synthesis of both G and A nucleotides (Extended Data Fig. 2c), resulting in depletion of A nucleotides. Thus, exogenous G disrupts purine balance by increasing levels of G nucleotides while preventing the synthesis of A nucleotides.

To assess whether an analogous imbalance in relative nucleotide levels can account for impaired proliferation upon salvage of other individual nucleotide precursors (Fig. 1b-f), we compared intracellular nucleotide levels upon addition of A-, dA-, G-, dG-, T- and C- precursors, each at concentrations that inhibit proliferation. Each salvage precursor altered relative nucleotide levels in different ways (Fig. 2d); therefore a change in any specific nucleotide species does not explain decreased proliferation across these conditions. Rather, these results argue that cells are vulnerable to multiple different perturbations in the balance of nucleotide species. Adding nucleotide precursors at lower concentrations that do not inhibit proliferation had less effect on intracellular nucleotide levels (Extended Data Fig. 2d), further suggesting that individual nucleotides impair proliferation due to imbalanced nucleotide species. Indeed, providing adenine to reestablish purine balance restored proliferation of guanine-treated cells (Fig. 2e and Extended Data Fig. 2e). Providing cytidine to balance pyrimidine levels also restored proliferation of cells treated with thymidine (Fig. 2f). Together, these data suggest that both pyrimidine and purine imbalance can inhibit proliferation.

In addition to perturbing ribonucleotide balance, salvage of individual nucleotides also altered intracellular dNTP levels. dGTP has the same molecular weight as ATP and similar chromatographic properties, and because ATP is much more abundant in cells, the peak for dGTP could not be confidently distinguished by LCMS. Nevertheless, we found that addition of each nucleotide precursor at concentrations that impair proliferation caused relative imbalances among dNTP species (Fig. 2g). Adding precursors at concentrations that do not affect proliferation had much less of an effect on dNTPs (Extended Data Fig. 2f), indicating that imbalanced dNTPs may also play a role in impairing proliferation upon nucleotide precursor addition.

### Nucleotide imbalance slows progression through S phase of the cell cycle

The association between altered dNTP levels and decreased proliferation raised the possibility that nucleotide imbalance impairs proliferation by impeding DNA replication during S phase. To test this, we monitored cell cycle progression following G nucleotide treatment using flow cytometry to measure both DNA content and incorporation of EdU into DNA, which reflects active DNA replication during S phase (Fig. 3a). Untreated, asynchronous cells contain populations in G1 phase (with 2N DNA content), S phase (with intermediate DNA content and EdU incorporation), and G2/M phases (with 4N DNA content). As reported in numerous classic studies, serum starvation causes cells to accumulate in G1, while Taxol treatment causes cells to accumulate in G2/M^26^ (Extended Data Fig. 3a). In line with its ability to inhibit proliferation (Fig. 1b), guanine treatment had a dose-dependent effect on cell cycle progression. Addition of guanine at increasing concentrations caused cells to accumulate in S phase by 24 hours, and at the highest concentration cells failed to robustly incorporate EdU (Fig. 3b and Extended Data Fig.3b). Similarly, providing guanine at a concentration that inhibits proliferation over 96 hours initially increased the population of cells in S phase, and then prevented EdU incorporation (Extended Data Fig. 3c). Consistent with restoring nucleotide balance and rescuing proliferation, providing adenine together with guanine restored normal cell cycle distribution (Fig. 3c and Extended Data Fig. 3b). Altering pyrimidine balance by providing thymidine also caused cells to accumulate in S phase (Extended Data Fig. 3d). Treatment with LTX or BRQ to decrease total purine or pyrimidine levels, respectively, similarly prevented EdU incorporation by 96 hours. However, LTX or BRQ did not cause the same pattern of S phase accumulation as observed with G treatment (Extended Data Fig. 3c). This could suggest that the while purine or pyrimidine imbalance impairs proliferation by slowing S phase progression, overall purine or pyrimidine depletion may inhibit proliferation at least in part through a different mechanism.

**Figure 3.**
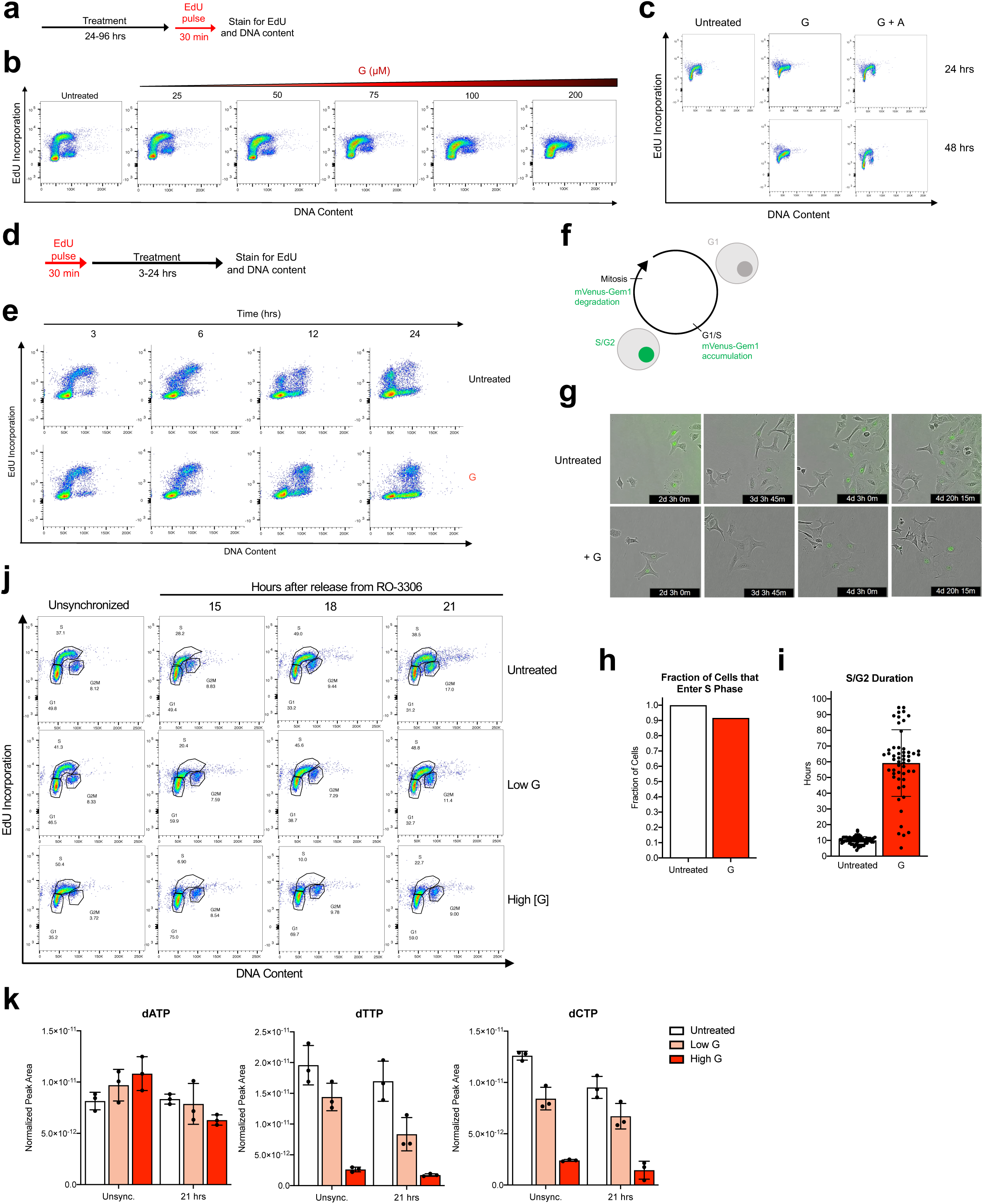
S phase progression is impaired by nucleotide imbalance. **a**, Experimental approach to assess cell cycle phase by DNA content and incorporation of EdU into DNA undergoing replication. DNA content (as determined by propidium iodide staining) and EdU incorporation were assessed by flow cytometry. **b**, Cell cycle distributions of A549 cells cultured in standard conditions or treated with the indicated concentration of guanine (G) for 24 hours. **c**, Cell cycle distributions of A549 cells treated with 200 µM G with or without 200 µM adenine (A) for 24 or 48 hours. **d**, Experimental approach to assess S phase progression. An initial pulse of EdU marks cells that are in S phase and therefore actively replicating DNA. EdU-positive and -negative cell populations are monitored as they progress though the cell cycle by measuring DNA content. **e**, Cell cycle distributions of A549 cells pulsed with EdU, and then cultured in standard conditions (Untreated) or treated with 200 µM G for the indicated amount of time. **f**, Schematic of fluorescent reporter system using expression of mVenus-Gem1 to assess cell cycle dynamics in live cells. **g**, Representative images from live-cell imaging of A549 cells expressing mVenus-Gem1 cultured in standard conditions (Untreated) or with 200 µM G, showing that cells enter S phase and have prolonged S/G2 duration under G treatment. (See also Supplementary Videos 1 and 3.) **h**, Fraction of cells cultured in standard conditions (Untreated) or with 200 µM G that began the experiment in G1 phase and entered S phase, as assessed by live-cell imaging of A549 cells expressing mVenus-Gem1. 76 cells were analyzed. **i**, Duration of S/G2 phase in cells cultured in standard conditions (Untreated) or with 200 µM G, as assessed by live-cell imaging and tracking of A549 cells expressing mVenus-Gem1. 115 cells were analyzed. **j**, Cell cycle distribution of A549 cells synchronized by treating with 4.5 µM RO-3306 for 18 hours to arrest cells in G2 phase, then removing RO-3306 to release cells from arrest for the indicated amount of time. At the time of release, cells were treated with standard culture media (Untreated), 25 µM G (low G), or 200 µM G (high G) as indicated. Cells were pulsed with EdU for 30 minutes prior to analysis at each time point as outlined in Fig. 3a. **k**, Intracellular levels of dNTPs in synchronized A549 cells 21 hours after release from RO-3306. At the time of release from RO-3306, cells were cultured in standard conditions (Untreated) or treated with 25 µM G (low G), or 200 µM G (high G) as indicated. Levels of dNTPs in unsynchronized cells cultured in untreated media or treated for 24 hours with low G or high G are also shown. dNTPs were measured using LCMS. Data are presented as mean +/- SD.

The lack of EdU incorporation at higher nucleotide concentrations could either indicate that cells do not enter S phase, or that they enter S phase but fail to incorporate EdU because they are unable to undergo continuous DNA replication. To more directly examine whether DNA replication is slowed following purine imbalance, we monitored the progression of specific cell populations through S phase. We first pulsed untreated cycling cells with EdU to mark the population of cells that are in S phase at t=0, then measured cell cycle progression with and without guanine supplementation over 24 hours. In an untreated sample, the EdU-positive population progressed to a 4N DNA content and then back to a 2N DNA content, reflecting completion of S phase and return to G1 after cell division (Fig. 3d and Extended Data Fig. 3e). While untreated EdU-positive cells completed S phase and divided within 24 hours, guanine-treated EdU-positive cells failed to divide by 24 hours (Fig. 3e), suggesting that S phase progression is slowed following induction of purine imbalance. Further, a population of EdU-negative cells with intermediate DNA content accumulated during the 24 hours of guanine treatment. Initially, EdU-negative cells with 2N DNA content are in G1 phase. Thus, accumulation of EdU-negative cells at intermediate DNA content argues that following induction of purine imbalance, cells enter S phase but progression through S phase is impaired.

To further examine how nucleotide imbalance affects the kinetics of S phase entry and duration, we performed live cell imaging using a previously described fluorescent reporter for cell cycle state^27^. Specifically, we used mVenus conjugated to a truncated form of geminin, whose degradation is cell cycle-dependent such that cells expressing mVenus-Gem1 have fluorescent nuclei between the G1/S transition and mitosis, allowing for specific monitoring of S phase entry and quantification of S/G2 and G1 durations (Fig. 3f,g and Supplementary Videos 1-3). Almost all guanine-treated cells entered S phase, but subsequently had drastically longer S/G2 duration than untreated cells (Fig. 3h,i). Guanine treatment also increased G1 duration in cells born after induction of nucleotide imbalance, though not to the same extent as S/G2 duration (Extended Data Fig. 3f).

Because salvage of individual nucleotides caused dNTP levels to become imbalanced in cell populations (Fig. 2g), we tested whether dNTP imbalance persisted as cells entered S phase by synchronizing cells in G2 phase using the CDK1 inhibitor RO-3306^28^ and then releasing cells into the following cell cycle. Importantly, this strategy does not directly perturb cell metabolism, allowing us to assess cell cycle-dependent metabolic states. Untreated cells entered S phase around 15 hours after release and progressed to late S phase by around 21 hours after release, while cells treated with high concentrations of guanine had slower S phase progression (Fig. 3j). Increasing concentrations of guanine caused increased dNTP imbalance 21 hours after release from RO-3306 (Fig. 3k), demonstrating that nucleotide imbalance perturbs dNTP availability during S phase. Together, these data show that cells do not have a mechanism to prevent them from entering S phase with imbalanced nucleotide levels. Instead, nucleotide imbalance results in impaired S phase progression and potential problems with DNA replication.

### Growth control pathways do not prevent biomass production under nucleotide imbalance, leading to continued cell growth without cell division

Upon metabolic stress, numerous signaling pathways decrease biomass production in order to prioritize survival and recovery of homeostasis^1^. As nucleotide metabolism is regulated by these pathways and supports both cell growth and division, we investigated whether signaling mechanisms decrease growth in coordination with decreased proliferation rate under nucleotide imbalance. In particular, mTORC1 coordinates biomass production, regulates nucleotide synthesis, and senses levels of specific nucleotides: depletion of all purines, but not pyrimidines, has been shown to inhibit mTORC1 signaling^11, 12^. However, we found that mTORC1 signaling remains active despite nucleotide imbalance (Fig. 4a and Extended Data Fig. 4a,d). Changes in the activity of other major growth regulatory pathways, Akt and AMPK, did not correlate with proliferation arrest caused by nucleotide imbalance (Extended Data Fig. 4b,c). Further, decreased mTORC1 activity upon nucleotide depletion did not prevent growth in some cells (Extended Data Fig. 4d,j), suggesting that metabolic signaling does not directly control growth in response to perturbed nucleotide homeostasis.

**Figure 4.**
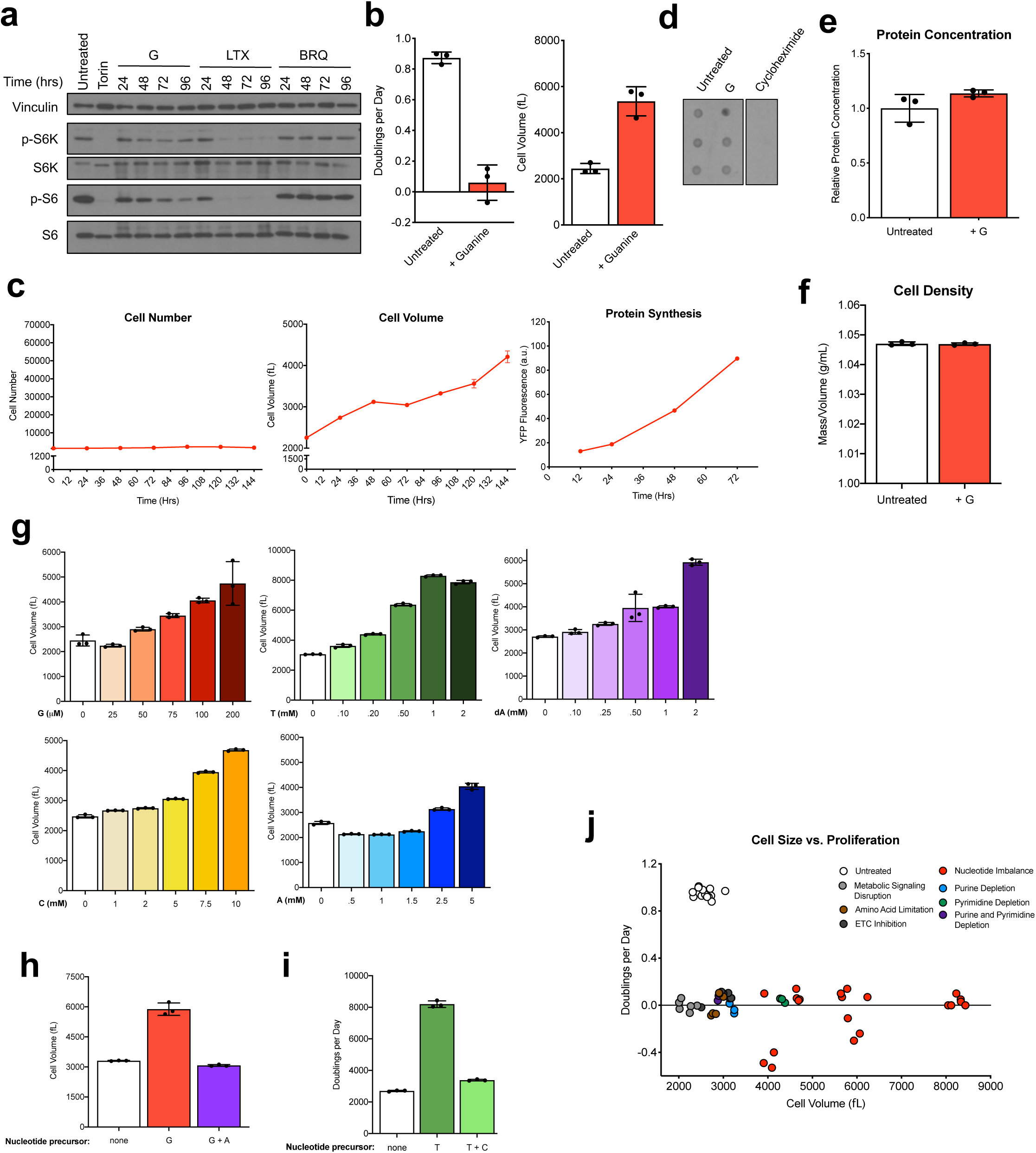
Imbalanced nucleotide levels lead to continued cell growth without division. **a**, Western blot showing phosphorylation of ribosomal protein S6 and S6 kinase (S6K) in A549 cells in standard culture media (Untreated) or following treatment with 1 µM Torin 1, or 200 µM guanine (G), 1 µM lometrexol (LTX), or 1 µM brequinar (BRQ) for the indicated amount of time. Levels of vinculin, total S6K, and total S6 were also determined as controls. **b**, Proliferation rate (left) and mean volume (right) of A549 cells cultured in standard conditions (Untreated) or treated with 200 µM guanine. **c**, Cell number (left), mean volume (center), and accumulation of newly synthesized protein (right) over time in A549 cells following treatment with 200 µM guanine. Protein accumulation was determined by quantifying fluorescence from A549 cells expressing YFP fused to a DHFR degron, which was stabilized by adding the ligand trimethoprim at t=0. **d**, Global protein synthesis as measured by incorporation of puromycin into nascent polypeptide chains. Puromycin was pulsed for 60 seconds in A549 cells cultured in standard conditions (Untreated) or treated for 24 hours with 200 µM G. Cycloheximide treatment was used as a negative control. **e**, Protein concentration in A549 cells cultured in standard conditions (Untreated) or treated with 200 µM G, calculated by dividing total protein measured per cell by the measured cell volume. **f**, Density of A549 cells cultured in standard conditions (Untreated) or with 200 µM G for 72 hours, calculated by dividing cell mass by cell volume. **g**, Mean volume of A549 cells treated for 96 hours with the indicated concentrations of G, thymidine (T), deoxyadenosine (dA), cytidine (C), or adenine (A). **h**, Mean volume of A549 cells cultured in standard media (none) or treated with 200 µM G with or without 200 µM A for 96 hours as indicated. **i**, Mean volume of A549 cells cultured in standard media (none) or treated with 1 mM T with or without 1 mM C for 96 hours as indicated. **j**, Correlation of proliferation rate and size of A549 cells cultured in conditions that perturb cell metabolism. Data are compiled from experiments shown in Fig. 1a-f, 2e, 2f, and 4g-i, and conditions are grouped into those that disrupt signaling (Torin treatment or serum starvation), amino acid limitation (leucine or arginine starvation), ETC inhibition (electron transport chain inhibition: oligomycin or rotenone treatment), depletion of purines (using LTX) or pyrimidines (using BRQ), or nucleotide imbalance induced by nucleobase/nucleoside supplementation. Data are presented as mean +/- SD of 3 biological replicates.

We next examined whether nucleotide imbalance allows continued cell growth and biomass production. Indeed, guanine-treated cells grew aberrantly large (Fig. 4b). Measuring production of a YFP protein synthesis reporter^29, 30^ showed that guanine-treated cells accumulated newly synthesized protein in coordination with increasing cell volume (Fig. 4c and Extended Data Fig. 4e). Evaluating puromycin incorporation into protein further argues that rates of protein synthesis are unaffected by purine imbalance (Fig. 4d). Thus, protein concentration and overall cell density are maintained despite a larger cell size (Fig. 4e,f). Other nucleotide imbalances that impaired proliferation also caused cells to grow aberrantly large, and similar to the anti-proliferative effect, the increase in size was dose-titratable and observed in all cell types tested (Fig. 4g and Extended Data Fig. 4f). A549 cell proliferation was most sensitive to G nucleotide precursors at lower concentrations (Extended Data Fig. 1e), and consistent with this, adding other nucleobases/nucleosides at concentrations that do not affect proliferation also did not change cell size (Extended Data Fig. 4g). Reestablishing purine or pyrimidine balance restored the normal size of cells treated with G- or T-nucleotide precursors, respectively (Fig. 4h,i). Moreover, most other metabolic perturbations did not increase cell size to a large extent, with the exception of pyrimidine synthesis inhibition (Extended Data Fig. 4h). Thus, while metabolic state, growth, and proliferation are normally tightly linked, these data suggest that cell growth is decoupled from proliferation following nucleotide imbalance (Fig. 4j).

The purine synthesis inhibitor LTX depletes both A- and G-nucleotides and prevents growth of some but not all cells (Extended Data Fig. 1a, and Extended Data Fig. 4i,j). In cells where LTX prevents growth and inhibits mTORC1 signaling, we asked whether supplementing purine-depleted cells with either adenine or guanine alone could cause purine imbalance and therefore decouple growth from proliferation. Adenine and guanine have been shown to reactivate mTORC1 signaling in purine-depleted cells, but the time required for A- versus G- nucleotides to induce mTORC1 activity may be variable^11, 12^. We found that both guanine and adenine could activate mTORC1 acutely following purine depletion, as well as sustain signaling over a longer time period (Extended Data Fig. 4k,l). However, activation of growth signaling is not sufficient to support proliferation: providing excess adenine or guanine did not restore proliferation (Extended Data Fig. 4m). Of note, low concentrations of adenine that do not induce nucleotide imbalance could rescue proliferation of LTX-treated cells. This may be explained by the ability of AMP deaminase to convert AMP to IMP, which can be then be converted to GMP, and sufficiently replenish both A- and G- nucleotides. Together, these data suggest that while sufficient levels of either purine can restore growth, purine balance is required for proliferation. Further, while purine-depleted cells (with inactive mTORC1 signaling) accumulate in G1 phase of the cell cycle, providing guanine causes these cells to enter S phase but be unable to complete S phase (Extended Data Fig. 4n). We therefore hypothesized that mTORC1 activity is needed for S phase entry in cells with nucleotide imbalance, and consistent with this, pharmacological inhibition of mTORC1 prevented guanine-treated cells from entering S phase (Extended Data Fig. 4n).

### DNA damage and replication stress-sensing pathways are activated upon nucleotide imbalance

Impaired S phase progression suggests that nucleotide imbalance leads to stalled DNA replication, and we therefore tested whether cells exhibit evidence of replication stress under nucleotide imbalance. The ATR and ATM kinases are well-characterized components of the cellular response to DNA damage in S phase, and sense single-stranded DNA and DNA double-strand breaks, respectively^31–33^. Chk1 and Chk2 are the respective targets of ATR and ATM, and are major effectors of the DNA damage response (Fig. 5a). Guanine treatment caused robust phosphorylation of both Chk1 and Chk2, with higher concentrations of guanine that inhibit proliferation to a greater extent inducing a stronger signaling response (Fig. 5b). Interestingly, the appearance of p-Chk1 occurred first by 24 hours of guanine treatment, followed by the appearance of p-Chk2 between 48 and 72 hours. This may indicate that replication fork stalling occurs first following purine imbalance and activates ATR, with later activation of ATM. Addition of adenine together with guanine prevented induction of the replication stress response (Extended Data Fig. 5a). Nucleotide imbalances induced by other precursors also activated ATR and ATM signaling, while as expected, using leucine deprivation to inhibit proliferation did not (Fig. 5c and Extended Data Fig. 5a). Together, this suggests that impaired DNA replication is a reason why nucleotide imbalance prevents proliferation. Inhibiting total purine or pyrimidine synthesis induced phosphorylation of Chk1 and Chk2 to a lesser extent than guanine treatment, consistent with fewer cells stalling in S phase in these conditions (Fig. 5d and Extended Data Fig. 3b).

**Figure 5.**
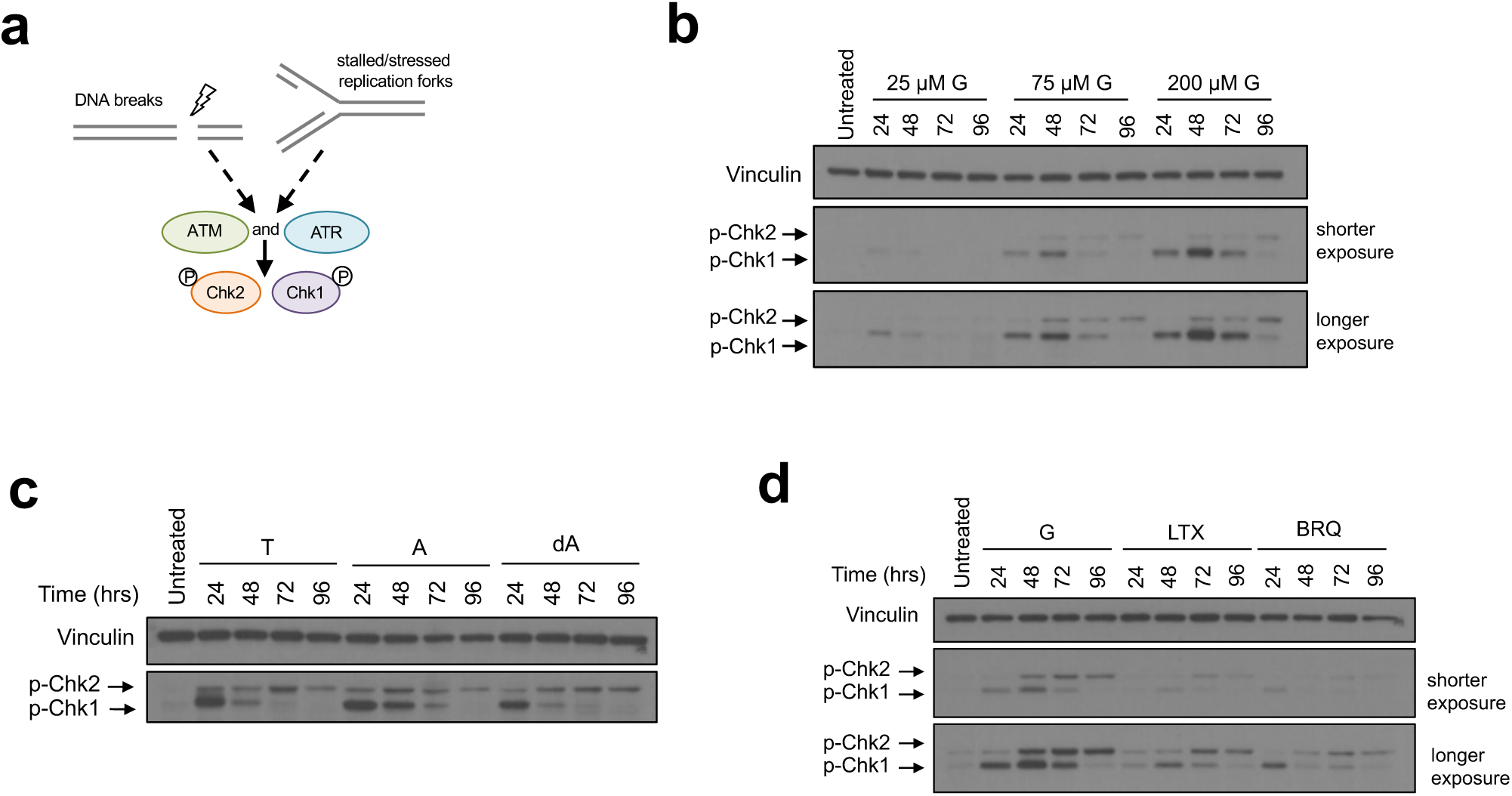
Nucleotide imbalance induces a replication stress response. **a**, Schematic outlining how ATR and ATM kinases respond to replication stress and DNA damage. ATR and ATM phosphorylate Chk1 and Chk2, respectively. **b**, Western blot showing phosphorylation of Chk1 and Chk2 in A549 cells cultured in standard conditions (Untreated) or treated for the indicated time with the indicated concentration of guanine (G). **c**, Western blot showing phosphorylation of Chk1 and Chk2 in A549 cells cultured in standard conditions (Untreated) or treated for the indicated amount of time with 1 mM thymidine (T), 2.5 mM adenine (A), or 1.5 mM deoxyadenosine (dA). **d**, Western blot showing phosphorylation of Chk1 and Chk2 in A549 cells cultured in standard conditions (Untreated) or treated for the indicated time with 200 µM G, 1 µM lometrexol (LTX), or 1 µM brequinar (BRQ). Levels of vinculin are also shown in all Western blots as a loading control.

Recent work demonstrated that pharmacologically inhibiting G nucleotide synthesis using the IMPDH inhibitor mycophenolic acid (MPA) can have dose-dependent effects. Treating cells with a low dose of MPA for 24 hours increased p53 and p21 protein levels and caused cells to accumulate in G1 phase, whereas a high dose of MPA caused p21 degradation and increased the number of cells in S phase^34^. We tested whether excess guanine addition has similar dose-dependent effects on p53 and p21 protein levels. In line with its effects on ATR and ATM signaling, increasing concentrations of guanine lead to increased p53 levels; however, higher guanine concentrations did not increase p21 degradation (Extended Data Fig. 5b). This suggests that inhibiting nucleotide synthesis and excess nucleotide salvage have differing effects on cells.

The role of ATR and ATM in the cellular response to DNA damage has been extensively studied^31, 32^; however, only a small fraction of guanine-treated cells exhibited relatively minor increases in DNA damage at 24 hours as measured by a comet tail assay (Extended Data Fig. 5c). At that time, the signaling response is already robust, indicating that replication stress-sensing pathways are activated under nucleotide imbalance without large amounts of DNA damage. Further, the finding that metabolic regulatory mechanisms fail to halt cell growth and prevent S phase entry downstream of nucleotide imbalance suggests that replication stress sensing constitutes the major signaling response to nucleotide imbalance.

### Replication stress signaling is required for cells to survive and recover from nucleotide imbalance

ATR and ATM activate downstream effectors that block cell cycle progression to prevent cell division (Extended Data Fig. 6a)^31, 33, 35^. ATR-mediated cell cycle arrest might therefore explain why nucleotide imbalance prevents proliferation. If this is the case, inhibiting ATR signaling would be expected to allow cells to continue proliferating despite nucleotide imbalance. After confirming that the ATR inhibitor AZ20^36, 37^ blocks downstream signaling (Extended Data Fig. 6b), we tested whether inhibiting ATR allowed for cell proliferation under purine imbalance. Instead of restoring proliferation, ATR inhibition sensitized cells to guanine treatment, increasing cell death (Fig. 6a and Extended Data Fig. 6c,d). ATR inhibition increased sensitivity to all nucleotide imbalances (Fig. 6b and Extended Data Fig. 6e), indicating that rather than being detrimental for proliferation, ATR activity can be a protective mechanism to allow cell survival with nucleotide imbalance. ATR inhibition did not sensitize cells to purine or pyrimidine depletion (Extended Data Fig. 6f), consistent with a less robust induction of ATR signaling in these conditions. This suggests that the replication stress response enables cell adaptation to nucleotide imbalance.

**Figure 6.**
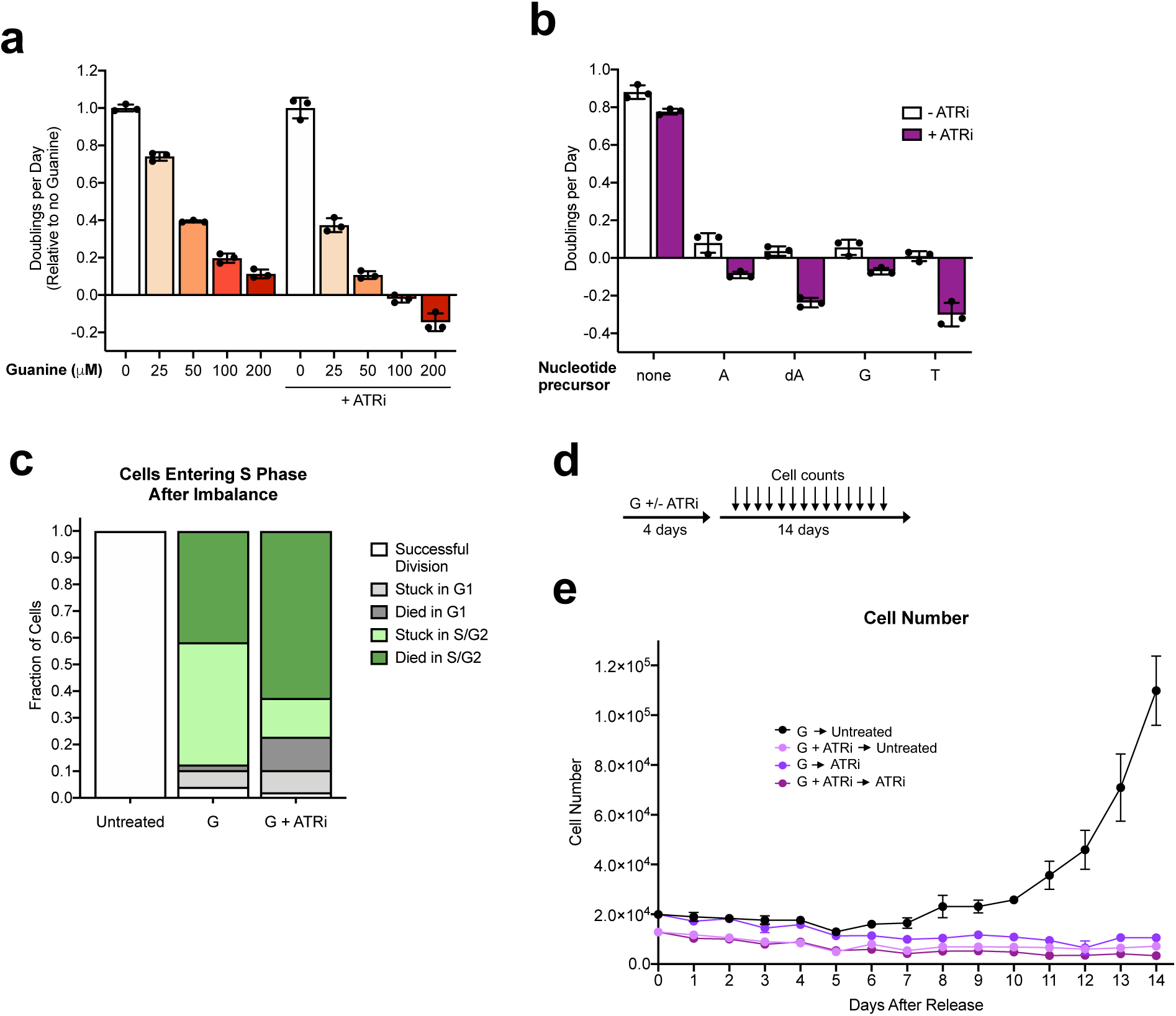
Replication stress signaling under nucleotide imbalance promotes cell survival and enables proliferation when nucleotide balance is restored. **a**, Proliferation rates of A549 cells treated with the indicated concentration of guanine with or without 50 nM of the ATR kinase inhibitor AZ20 (ATRi) as indicated. b, Proliferation rate of A549 cells cultured in standard conditions (none) or treated with 2 mM adenine (A), 1.5 mM deoxyadenosine (dA), 200 µM guanine (G), or 1 mM thymidine (T), with or without 50 nM ATRi as indicated. c, Cell fate of A549 cells expressing the mVenus-Gem1 reporter that were in G1 phase at the time of addition of 200 µM G with or without 50 nM ATRi, as assessed using live-cell imaging. The fate of cells in that were in G1 at the beginning of the experiment and were not exposed to excess G is also shown (Untreated). 124 cells were analyzed. d, Schematic showing the experimental approach to assess how cells recover from treatment with excess G, with or without inhibition of ATR signaling (ATRi). For these experiments, cells were cultured in media containing 200 µM G with or without 50 nM ATRi for 4 days. Media was then changed to untreated media or media containing 50 nM ATRi and cell number was determined every 24 hours for 14 days thereafter. e, A549 cell number over time after release from treatment with G with or without ATRi treatment as outlined in d. Data are presented as mean +/- SD of 3 biological replicates.

Live imaging with the mVenus-Gem1 cell cycle reporter (Fig. 3f) revealed that cells can successfully divide if they have partially completed S phase before guanine is added to the media, though ATR inhibition caused some of these cells to arrest or die (Extended Data Fig. 6g). This is consistent with the expected kinetics of nucleotide imbalance, as these cells turn over purine nucleotide pools in approximately 24 hours^17^. Thus, salvage of excess guanine and subsequent inhibition of A nucleotide synthesis (Fig. 2c and Extended Data Fig. 2c) would not be expected to immediately deplete A nucleotides. Cells that have already partially replicated their DNA likely can complete S phase before the balance of A and G nucleotides is drastically changed. In contrast, cells that are in G1 at the time of guanine supplementation and enter S phase with imbalanced nucleotides are unable to divide, and instead either arrest or die after entering S phase (Fig. 6c). Inhibiting ATR signaling caused most cells to die when they entered S phase with imbalanced purines, suggesting that replication stress sensing is critical for preventing S phase catastrophe under nucleotide imbalance. Interestingly, daughter cells born later after induction of nucleotide imbalance and ATR inhibition were more likely to become stalled in G1 phase (Extended Data Fig. 6h). Replication stress in mother cells can affect G1 length and lead to quiescence in daughter cells^38^, potentially implying that an inadequate replication stress response under nucleotide imbalance can result in DNA damage that is inherited by daughter cells and affects their proliferative potential.

We next asked whether cells can recover from purine imbalance caused by guanine supplementation, and found that when the imbalance is removed, cells resume proliferation at a normal size. However, ATR inhibition either concurrent with or after guanine treatment prevented recovery (Fig. 6d,e, and Extended Data Fig. 6i). This further argues that failure to activate a replication stress response under nucleotide imbalance causes irreversible damage to cells. ATR inhibition can allow inappropriate firing of late origins of DNA replication, and we reasoned that this may force cells to continue through S phase at a pace that is incompatible with relative dNTP availability. However, ATR inhibition did not appear to accelerate cell cycle progression upon guanine treatment (Extended Data Fig. 6j), suggesting that specific dNTPs themselves may become limiting as substrates for replication following nucleotide imbalance.

### Activation of the replication stress response supports increased dNTP availability during unperturbed S phases

That cells enter S phase with imbalanced nucleotides argues that cells do not monitor relative nucleotide levels prior to S phase entry, and replication stress signaling may be important for adjusting nucleotide availability during normal proliferation. Indeed, ATR activity has been observed in unperturbed cell cycles^35, 39, 40^, and the yeast ATR homolog Mec1 is activated as a result of low dNTPs at the onset of S phase^4^. To test whether replication stress signaling allows proliferating mammalian cells to maintain sufficient dNTPs for DNA replication, we first monitored replication stress sensing during normal S phases by synchronizing cells in G2 phase using the CDK1 inhibitor RO-3306^28^ and releasing cells into the following cell cycle. Chk1 phosphorylation, reflecting ATR activity, was induced as most cells entered early S phase, and was attenuated as most cells progressed through late S phase (Fig. 7a,b and Extended Data Fig. 7a).

**Figure 7.**
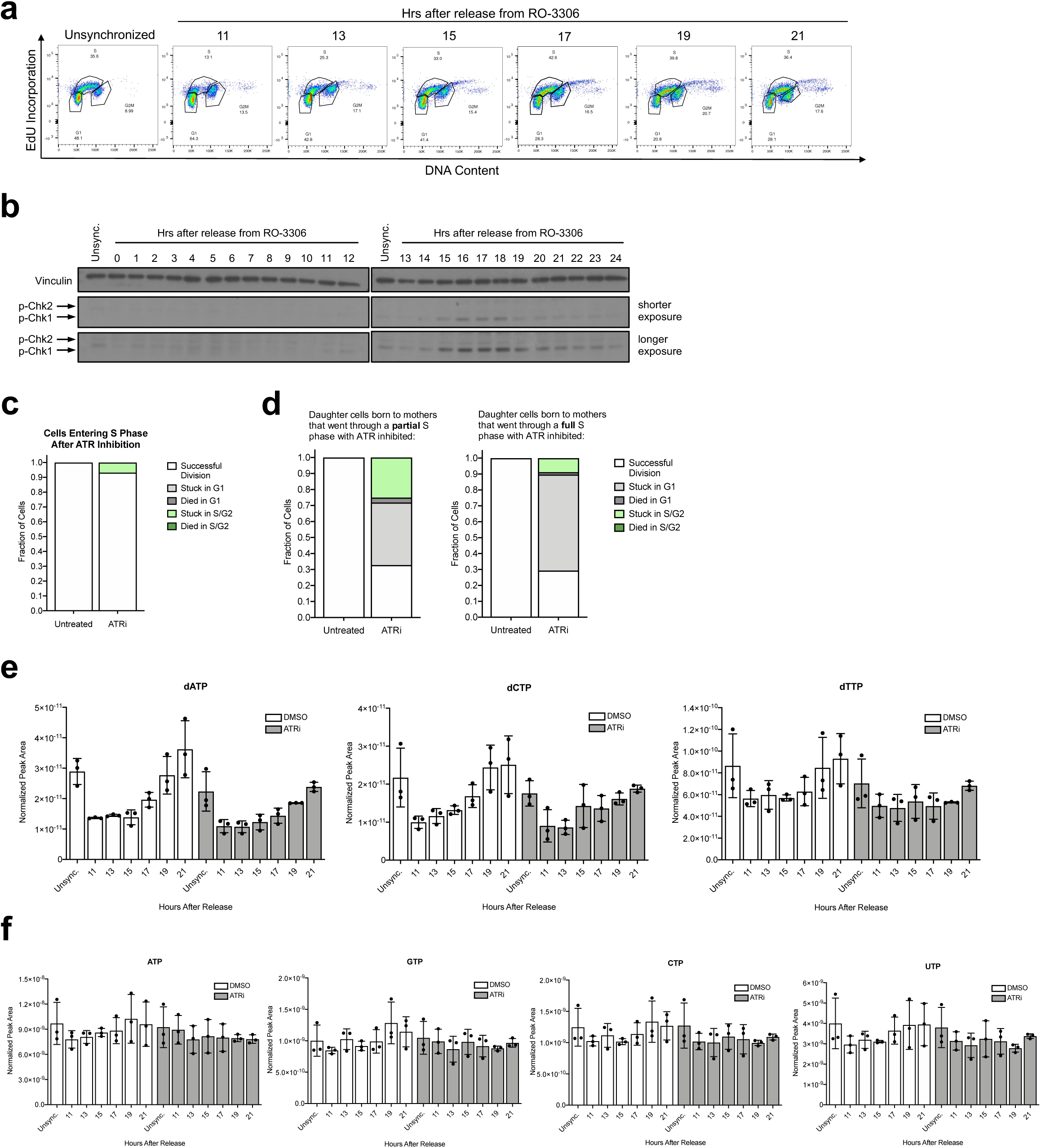
The replication stress response promotes increased dNTP levels during unperturbed S phase. **a**, Cell cycle distribution of A549 cells corresponding to the Western blots shown in panel **b**. Cells were treated with 9 µM RO-3306 for 18 hours to arrest cells in G2 phase, then RO-3306 was removed to release cells from arrest for the indicated amount of time. Cells were pulsed with EdU for 30 minutes prior to analysis at each time point as outlined in Fig. 3a. **b**, Western blot showing phosphorylation of Chk1 and Chk2 in unsynchronized A549 cells (Unsync.) or A549 cells treated with 9 µM RO-3306 for 18 hours to arrest cells in G2 phase, then released into the cell cycle for the indicated amount of time. Levels of vinculin are also shown as a loading control. **c**, Cell fate as assessed using live-cell imaging of A549 mother cells expressing the mVenus-Gem1 reporter that were in G1 phase at the time of 50 nM AZ20 (ATRi) addition. The fate of mother cells in G1 not exposed to ATRi is also shown (Untreated). 55 cells were analyzed. **d**, Cell fate of A549 daughter cells expressing the mVenus-Gem1 reporter born to mother cells either in S/G2 phase (left) or G1 phase (right) at the time of 50 nM ATRi addition. Mother cells that were in S/G2 phase when ATRi was added went through a partial S phase with ATR inhibited, whereas mother cells that were in G1 phase went through a full S phase with ATR inhibited. The fate of daughter cells not exposed to ATRi is also shown (Untreated). For the left and right panels, 104 and 107 cells were analyzed, respectively. **e**, dNTP levels in A549 cells synchronized in G2 phase by treating with 4.5 µM RO-3306 for 18 hours, then released from RO-3306 for the indicated times. At the time of release from RO-3306, cells were either treated with DMSO or 50 nM ATRi as indicated. Unsynchronized cells (Unsync.) were treated with DMSO or 50nM ATRi for 24 hours as indicated. **f**, Ribonucleotide levels in A549 cells synchronized in G2 phase by treating with 4.5 µM RO-3306 for 18 hours, then released into the cell cycle for the indicated amount of time. At the time of release from RO-3306, cells were either treated with DMSO or 50 nM ATRi as indicated. Unsynchronized cells (Unsync.) were treated with DMSO or 50 nM ATRi for 24 hours as indicated. All nucleotide levels were measured using LCMS. Data are presented as mean +/- SD of 3 biological replicates.

To explore whether ATR signaling is important for normal proliferation, we monitored how cell cycle progression and cell fate is affected by inhibiting ATR activity in mVenus-Gem1-expressing cells. On a population level, both G1 and S/G2 duration increased upon ATR inhibition, with G1 duration increasing to a greater extent (Extended Data Fig. 7b). Mother cells treated with ATR inhibitor successfully completed their current cell cycle (Fig. 7c and Extended Data Fig. 7c). However, the majority of daughter cells born to ATR-inhibited mother cells did not divide in a normal timeframe, and the plurality of daughter cells became stalled in G1 (Fig. 7d). The likelihood of G1 stalling was greater for daughter cells whose mothers underwent an entire S phase with inhibited ATR activity, compared to daughters whose mothers only experienced ATR inhibition for the latter part of S phase. These results are consistent with the idea that an inability to activate ATR signaling during an otherwise unperturbed S phase may result in DNA damage that is inherited by daughter cells and causes them to stall in G1^38^.

We next asked whether ATR activity had an effect on dNTP levels in cells progressing through S phase. While levels of other metabolites, including amino acids and ribonucleotides, were relatively constant throughout the cell cycle, dNTP levels were lower at a time point when the majority of cells entered S phase and increased as cells progressed through S phase (Fig. 7e,f and Extended Data Fig. 7d,e). Thus, ATR activation correlates with low dNTP levels upon S phase entry. These data are consistent with cells entering S phase with insufficient dNTPs for replication fork progression, leading to replication stress. Because downstream effectors of ATR can activate nucleotide synthesis enzymes^41^, we asked whether ATR activity helps cells meet this dNTP demand during normal proliferation. Indeed, ATR inhibition attenuated the increase in dNTP levels observed over the course of S phase (Fig. 7e), suggesting that ATR activity helps maintain dNTPs during normal S phases.

Together, these data suggest that cells do not sense nucleotide levels in preparation for S phase, but instead rely on replication stress signaling to modulate dNTP availability for genome replication.

## Discussion

In order to proliferate, cells must coordinate growth signaling with sufficient metabolite availability. Complex regulatory pathways integrate this information, allowing cells to balance anabolic and catabolic reactions in response to available nutrients. Under stress, a key response of these pathways is to halt biomass production in favor of prioritizing resources for survival until conditions improve. Consistent with this logic, purine depletion has been shown to inactivate growth signaling via inhibiting mTORC1^11, 12^. Surprisingly, a disruption in the balance of purine and pyrimidine nucleotides is not directly sensed, resulting in continued cell growth and protein production despite the inability proliferate.

Unexpectedly, the lack of mechanism to sense relative purine and pyrimidine balance results in cells entering S phase and initiating DNA replication despite the imbalance. Ultimately this means that imbalance leads to DNA replication stress and triggers checkpoints to respond to this stress. The fact that cells continue to produce protein argues that RNA synthesis is largely not impaired by nucleotide imbalance, despite ribosomal RNA accounting for the majority of nucleic acid biomass in cells. One possibility is that even when their levels are imbalanced, the availability of each ribonucleotide species is still sufficient for RNA synthesis. For example, while guanine treatment decreased intracellular adenylate nucleotide pools, baseline ATP levels are high relative to dNTPs, and may be unlikely to become limiting for RNA production.

The ATR and ATM signaling pathways are well-known mediators of the cellular response to DNA damage, but are activated under nucleotide imbalance even in the absence of extensive DNA damage. In fact, ATR and ATM signaling are critical for survival under conditions of nucleotide imbalance, potentially because these signals activate enzymes to stabilize replication forks and replenish dNTPs while preventing the firing of additional origins of replication. Recent evidence shows that ATR is activated and important for the proper sequence of origin firing during unperturbed S phases^35, 39, 40^. Further, ATR effectors can activate RNR to promote dNTP synthesis^41, 42^. Our finding that ATR activity is needed to increase dNTP availability during unperturbed S phases is consistent with a role for replication stress signaling in responding to nucleotide levels, and suggests that ATR may be important for allowing cells to adapt to fluctuating nucleotide levels encountered during normal cell divisions. In budding yeast, Mec1 (the yeast ATR homolog) is activated in early S phase downstream of initially low dNTP pools^4^, indicating that this metabolic role may be conserved across eukaryotic species.

ATR signaling during normal S phase could reflect reduced levels of all dNTPs or imbalanced dNTPs. Given that dNTPs are maintained at different relative concentrations and are rapidly consumed during genome replication, it is possible that dNTP imbalance occurs stochastically in a given cell as levels of one dNTP species decease more than others. A stochastic decrease in different dNTPs in different cells would also lead to reduced levels of all dNTPs being measured in a population. Regardless, our findings suggest that cells cannot sense whether they have the right levels of each nucleotide to meet the demands of replication before they enter S phase, and instead sense the downstream consequences of insufficient dNTPs and nucleotide imbalance.

Both salvage and synthesis pathways control cellular levels of purines and pyrimidines, and environmental conditions can differentially impact the use of these pathways to produce specific nucleotide species. Levels of available nucleotide salvage precursors vary depending on tissue context, dietary nutrients, and effects of other cells in the microenvironment^14, 43, 44^. As nucleotide synthesis incorporates metabolites from many aspects of metabolism, these same environmental factors could also differentially impact substrate availability to produce different nucleotide species. Levels of different nucleotide species, as well as expression of synthesis pathways, salvage enzymes, and transporters are expected to vary based on cell type. This may be why sensitivity to supplementation with different nucleotide species varies both within and across cell lines.

Because imbalance is not sensed by pathways that would limit S phase entry, rapidly proliferating cancer cells may be especially vulnerable to nucleotide imbalance. Many cancers harbor mutations in DNA damage response (DDR) pathways^45^. As replication stress signaling is essential for survival and recovery from nucleotide imbalance, loss of function in these pathways could render DDR-deficient tumors sensitive to perturbed nucleotide balance. Dysregulated expression of nucleotide salvage and catabolism enzymes may also render certain cancers vulnerable to imbalance. Recent work has shown that the dNTP-degrading enzyme SAMHD1 protects against cytotoxic dGTP buildup upon deoxyguanosine supplementation. SAMHD1-deficient tumor cells are sensitive to dGTP accumulation caused by deoxyguanosine supplementation and purine nucleoside phosphorylase (PNP) inhibition^46, 47^. Nucleotide imbalance is also implicated in non-cancer human disease settings. In particular, PNP deficiency and adenosine deaminase (ADA) deficiency are marked by aberrant accumulation of dGTP and dATP, respectively^48–50^. These disorders lead to severe immunodeficiencies due to insufficient T and B cell proliferation and ameliorating nucleotide imbalance may improve fitness of these cells.

While replication stress can result in irreversible growth arrest and senescence, we find cells can recover from impaired replication caused by nucleotide imbalance and resume proliferation, although the time needed to complete S phase can be long. Moreover, the observation that cells recover from imbalance suggests that senescence induction is not an early response to nucleotide imbalance in all cells. Nonetheless, replication stress is well documented to contribute to senescence^51, 52^, but this may require imbalances over extended time periods or a disruption of the downstream pathways that allow cells to recover from replication stress. In fact, in this study, cells treated with ATR inhibitor such that they do not recover from nucleotide imbalance continue to grow excessively large, consistent with classic descriptions of cellular senescence.

The observation that mean cell volume returns to normal upon recovery from nucleotide imbalance suggests that cells have an established “target” size and the cell population has the ability to return to that average size. Division is likely necessary for cell volume reduction, consistent with the observation that cell number begins to increase before mean size decreases. In addition, cell growth plateaus with prolonged arrest due to nucleotide imbalance, implying that mechanism(s) also exist to halt biomass production despite the initial uncoupling of cell growth and division. Nevertheless, it is unclear why cells continue to grow upon release from nucleotide imbalance, despite already being aberrantly large, suggesting that growth is not initially tightly controlled with respect to target cell size.

More generally, this study shows that cell growth and division can be uncoupled downstream of nucleotide imbalances that might occur in response to fluctuating nutrient levels. Replication stress signaling modulates nucleotide availability during normal proliferation and protects against fluctuations in nucleotide levels, but larger environmental changes that affect nucleotide balance increase the risk of genomic damage, raising the possibility that nucleotide imbalance-induced replication stress plays additional roles in cell physiology or function.

## Acknowledgements

We thank members of the Vander Heiden lab for helpful discussions, A. Darnell for constructs for the YFP-DHFR reporter, and K. Sapp for constructs for the mVenus-Gem1 reporter. The A9 cells were a gift from the B. Manning laboratory. We also thank the Koch Institute Flow Cytometry Core Facility. This research was supported by the National Cancer Institute of the NIH under award number F31CA236036 (F.F.D.) and award numbers R35CA242379, R01CA201276, and P30CA14051 (M.G.V.H.). T.P.M was supported by the Wellcome Trust grant 110275/Z/15/Z. M.G.V.H. also acknowledges support from a Faculty Scholar grant from the Howard Hughes Medical Institute, SU2C, a division of the Entertainment Industry Foundation, the Lustgarten Foundation, the MIT Center for Precision Cancer Medicine, and the Ludwig Center at MIT.

## Author Contributions

Conceptualization, F.F.D. and M.G.V.H.; Investigation, F.F.D., T.P.M., R.E., C.S.N., C.A.L.; Writing –Original Draft, F.F.D.; Writing – Review & Editing, F.F.D., T.P.M., R.E., C.S.N., S.R.M., C.A.L., M.G.V.H.; Supervision, M.G.V.H.; Funding Acquisition, F.F.D. and M.G.V.H.

## Declaration of Interests

The authors are aware of no direct conflicts with the topic of the study, however S.R.M. declares he is a co-founder of Affinity Biosensors and Travera, and M.G.V.H. declares he is a member of the scientific advisory board member for Agios Pharmaceuticals, Aeglea Biotherapeutics, Faeth Therapeutics, and iTeos Therapeutics, and a co-founder of Auron Therapeutics.

## Methods

### Cell Lines and Cell Culture

All cell lines were cultured in Dulbecco’s Modified Eagle’s Medium (DMEM) (GIBCO) supplemented with 10% heat inactivated fetal bovine serum at 37°C with 5% CO_2_. All cell lines regularly tested negative for mycoplasma. To generate cells with stable transgene expression, Lenti-X 293T cells at 75% confluency were transfected using X-tremeGENE 9 DNA transfection reagent (Sigma). The lentiviral plasmids used were pRSV-Rev (Addgene #12253), pMDLg/pRRE (Addgene #12251), and pMDG2.G (Addgene #12259) from Didier Trono. For the YFP-DHFR reporter, the donor expression plasmid pLJM1-FLAG-YFP-DHFR ^30^, was used. For the mVenus-Gem1 reporter, the donor expression plasmid pLenti-puro-mVenus-Gem1 was used. After 48 hours, lentivirus was harvested by removing the culture media from the Lenti-X 293T cells and passing it through a 0.45 μm filter. The target cell lines at 50-60% confluency were then infected using 3 mL virus with polybrene reagent (Sigma). After 24 hours, virus was removed and cells were allowed to recover in virus-free media for 24 hours. Selection was then initiated using puromycin at a concentration of 2 μg/mL.

### Proliferation Rates and Cell Size Measurements

All cell lines were plated in 6-well plates in DMEM with 10% FBS at a concentration of 20,000 cells per well with the exception of MDA-MB-468 cells, which were plated at a concentration of 40,000 cells per well. The number of cells seeded for each cell line allowed for exponential growth over the course of the assay. The following day, one 6-well plate of each cell line was counted to determine the initial number of cells at the time of treatment. Cells were washed three times with PBS and 4 mL of treatment media was added. Treatment media was made with 10% dialyzed FBS. Media lacking specific amino acids was made from DMEM without pyruvate or amino acids supplemented with an amino acid mix containing DMEM-concentrations of amino acids without arginine, leucine, or serine. Arginine, leucine, or serine were added back to the media as needed. After 4 days of treatment, final cell counts were measured using a Multisizer 3 Coulter Counter (Beckman Coulter). The following formula was used to calculate proliferation rate:

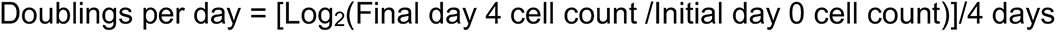

Cell size measurements were taken using a Multisizer 3 Coulter Counter at the same time as cell counts, after 4 days of treatment.

### Cell Density

The average cell density in a population was measured by comparing Multisizer 3 Coulter Counter (Beckman Coulter)-based cell volume measurements and suspended microchannel resonator (SMR)-based buoyant mass measurements. The resulting relative density values were converted to absolute density values by also measuring the density of the measurement solution (culture media) using SMR and by calibrating the SMR measurements using polystyrene beads of known volume^53, 54^. Population average densities were measured after 3 days of drug treatment, because longer drug treatments resulted in cells that were too large for the SMR microchannels.

### Protein Synthesis: Puromycin Incorporation

25,000 cells were plated in 6 cm plates. The following day, cells were washed three times with PBS and 5 mL of treatment media containing 10% dialyzed FBS was added for the specified amount of time. To perform the puromycin pulse, cells were kept at 37°C and puromycin was added to the culture medium at 10 μg/mL for exactly 1 minute. Cells were washed once in ice-cold PBS, and the plates were flash frozen in liquid nitrogen and subsequently stored at -80°C. 10 μg/mL cycloheximide was added to a negative control plate 30 minutes prior to the puromycin pulse. Protein lysates were prepared using ice-cold RIPA buffer with protease inhibitor. To perform dot blots, samples were normalized for protein concentration and 2 µL of lysate was dotted onto a 0.22 µm nitrocellulose membrane. Membranes were blocked with 5% milk for 1 hour, washed, and incubated at 4°C overnight with the following primary antibodies in 5% BSA in TBST: puromycin (Sigma, 1:25,000) and vinculin (Cell Signaling Technology, 1:1,000). The following day, membranes were washed three times with TBST on a rocker for 10 minutes. Secondary antibodies were applied for 60 minutes. Anti-rabbit (Cell Signaling Technology) secondary antibody was used at a dilution of 1:5000 and anti-mouse (Cell Signaling Technology) secondary antibody was used at a dilution of 1:10000. Membranes were then washed again three times with TBST for 10 minutes and signal was detected with ECL using film.

### Protein Synthesis: YFP Reporter

To assess protein production and accumulation over time, a previously described protein synthesis reporter was employed ^29^ ^30^. In this reporter system, YFP is fused to an engineered unstable *E. coli* dihydrofolate reductase (DHFR). YFP is rapidly degraded, and only accumulates in the presence of the ligand trimethoprim (TMP), which stabilizes the DHFR domain. Accumulation of fluorescence over time in the presence of TMP therefore reflects protein synthesis rate of the YFP reporter. To monitor YFP production, 80,000 cells were seeded in 6-well plates and allowed to adhere overnight. The following day, cells were washed three times with PBS and 4 mL of treatment media containing 10 µM TMP was added. After the indicated amount of time, cells were trypsinized, pelleted, and resuspended in PBS. YFP fluorescence was measured using flow cytometry.

### Protein Concentration

Protein concentration was calculated by dividing total protein content by cell number and cell volume for each sample. A BCA assay was used to measure total protein content, as compared to a standard curve. At the same time, cell number and volume were measured in a parallel sample using a Multisizer 3 Coulter Counter (Beckman Coulter).

### Cell Cycle Analysis by Flow Cytometry

500,000 cells were plated in 10 cm plates and incubated overnight to allow cells to adhere. The following day, cells were washed three times with PBS and 10 mL of treatment media with 10% dialyzed FBS was added for the desired amount of time. EdU was spiked into cell culture plates at 10 µM for exactly 30 minutes prior to fixing. To fix cells, each plate was trypsinized, pelleted and washed twice with PBS. Cells were resuspended in 500 µL ice-cold PBS and 5 mL ice-cold ethanol was added dropwise to each sample while vortexing in order to obtain a single-cell suspension. Fixed cells were stored at 4°C until being processed by flow cytometry (no longer than 4 days).

Cells were stained for EdU using the Click-iT EdU Pacific Blue kit (Invitrogen) according to manufacturer instructions. After staining for 30 minutes, cells were stained with propidium iodide. Cells were pelleted and washed with 1% BSA in PBS, then resuspended in 800 µL 1% BSA in PBS. 200 µL Propidium Iodide/RNAseA staining solution was added to each sample and cells were stained for at least 45 minutes at 4°C protected from light. Samples were then passed through a .35 µm filter into flow cytometry tubes (Falcon). Samples were run on a BD FACSCanto II Cell Analyzer, and 10,000 events were recorded for each sample.

### Comet Assay

500,000 cells were plated in 10 cm plates and incubated overnight to allow cells to adhere. The following day, cells were washed three times with PBS and 10 mL of treatment media containing 10% dialyzed FBS was added. After 24 hours of treatment, cells were trypsinized, pelleted, and resuspended in ice-cold PBS at a concentration of 1x10^5^ cells/mL. Sample preparation, electrophoresis, staining, and microscopy were then performed using a CometAssay Kit (Trevigen) according to manufacturer instructions. Percent of DNA in comet tail was quantified using ImageJ with OpenComet software^55^.

### Western Blots

10^6^ cells were plated in 10 cm plates and incubated overnight to allow cells to adhere. The following day, cells were washed three times with PBS and 10 mL of treatment media containing 10% dialyzed FBS was added. After culturing cells for the indicated time in treatment media, protein lysates were prepared by rapidly placing cells on ice and washing with ice-cold PBS, then lysing cells in ice-cold RIPA buffer containing cOmplete protease inhibitor (Roche) and phosphatase inhibitor cocktail (Sigma-Aldrich). Lysed cells were vortexed at 4°C for 10 min and then centrifuged at maximum speed at 4°C for 10 min. Protein lysate supernatant was removed and stored at -80°C. Proteins were separated using SDS-PAGE (12% acrylamide gels) and transferred to a nitrocellulose membrane using a standard wet transfer method. Membranes were blocked for 60 minutes using 5% bovine serum albumin (BSA) in Tris buffered saline with Tween (TBST). Membranes were incubated in primary antibody overnight at 4°C. The following primary antibodies were used at a dilution of 1:1000: vinculin (Cell Signaling Technology #4650), phospho-ribosomal protein S6 Ser 235/236 (CST #4858), ribosomal protein S6 (CST #2217), phospho-p70 S6 kinase Thr389 (CST #9205), p70 S6 kinase (CST #9202), phospho-Akt (CST #4060), Akt (CST #9272), phospho-AMPK (CST #2535), AMPK (CST #2532), phospho-Chk1 Ser345 (CST #2341), phospho-Chk2 Thr68 (CST,2197), p53 (CST #9282), and p21 (CST, #2947). Antibodies were diluted in 5% BSA in TBST. The following day, membranes were washed three times with TBST on a rocker for 10 minutes. Secondary antibodies were applied for 60 minutes. Anti-rabbit (CST #7074) secondary antibody was used at a dilution of 1:5000 and anti-mouse (CST #7076) was used a t a dilution of 1:10000. Membranes were then washed again three times with TBST for 10 minutes and signal was detected with ECL using film.

### LCMS Analysis

100,000 cells were plated in 6-well plates in DMEM with 10% FBS and incubated overnight. The following day, cells were washed three times with PBS and 4 mL of treatment media was added. All treatment media was made with 10% dialyzed FBS. After the indicated time period, polar metabolites were extracted from cells: plates were placed on ice, cells were washed with ice-cold blood bank saline, and 500 μl of ice-cold 80% methanol in water with 250nM ^13^C/^15^N labeled amino acid standards (MSK-A2-1.2: Cambridge Isotope Laboratories, Inc.) was added to each well. Cells were scraped, each sample was vortexed for 10 minutes at 4°C, and then centrifuged at maximum speed for 10 minutes at 4°C. Samples were dried under nitrogen gas and resuspended in 25 μl of a 50/50 acetonitrile/water mixture. Metabolites were measured using a Dionex UltiMate 3000 ultra-high performance liquid chromatography system connected to a Q Exactive benchtop Orbitrap mass spectrometer, equipped with an Ion Max source and a HESI II probe (Thermo Fisher Scientific). Samples were separated by chromatography by injecting 2-10 μl of sample on a SeQuant ZIC-pHILIC Polymeric column (2.1 × 150 mm 5 μM, EMD Millipore). Flow rate was set to 150 μl/min, temperatures were set to 25 °C for column compartment and 4 °C for autosampler sample tray. Mobile Phase A consisted of 20 mM ammonium carbonate, 0.1% ammonium hydroxide. Mobile Phase B was 100% acetonitrile. The mobile phase gradient (%B) was set in the following protocol: 0-20 min.: linear gradient from 80% to 20% B; 20-20.5 min.: linear gradient from 20% to 80% B; 20.5-28 min.: hold at 80% B. Mobile phase was introduced into the ionization source set to the following parameters: sheath gas = 40, auxiliary gas = 15, sweep gas = 1, spray voltage = −3.1kV, capillary temperature = 275 °C, S-lens RF level = 40, probe temperature = 350 °C. Metabolites were monitored in full-scan, polarity-switching, mode. An additional narrow range full-scan (220-700 m/z ) in negative mode only was included to enhance nucleotide detection. The resolution was set at 70,000, the AGC target at 1,000,000, and the maximum injection time at 20 msec. Relative quantitation of metabolites was performed with XCalibur QuanBrowser 2.2 (Thermo Fisher Scientific) using a 5 ppm mass tolerance and referencing an in-house retention time library of chemical standards. Metabolite measurements were normalized to the internal ^13^C/^15^N labeled amino acid standard and to cell number.

### Stable Isotope Tracing

100,000 cells were plated in 6-well plates in 2 mL DMEM with 10% FBS and incubated overnight. The following day, cells were washed three times with PBS and 4 mL of treatment media was added. All treatment media was made with 10% dialyzed FBS and 4 mM ^15^N-amide-glutamine. 200 µM ^13^C- guanine or ^13^C-adenine was added to the treatment media as indicated. Cells were cultured in treatment media for 24 hours before polar metabolites were extracted and analyzed as described in “LCMS Analysis.”

### Live-Cell Imaging with Cell Cycle Reporter

Live cell tracking of A549 cells expressing the mVenus-Gem1 reporter was carried out using the IncuCyte live cell imaging system (Sartorius). The cells were plated at 40,000 cells/well density on 6-well plates and the 4-day long chemical treatments were started 24 hours after plating. Imaging was performed every 45 min using 10X objective and 150 ms exposure time for the FITC channel. Cell tracking was carried out manually using the IncuCyte software. Only cells that were mVenus-Gem1 positive at some point during the 24-hour period prior to chemical treatment were tracked. Manual tracking recorded timings of cell cycle state change and/or cell death, as detected based on cell morphology. The tracking of each cell lineage lasted until the division of second cell generation or until the end of the 4-day drug treatment. If a cell remained arrested in a specific cell cycle state until the conclusion of the experiment, the duration of that cell cycle state was calculated to end at the conclusion of the experiment. Consequently, for a small fraction of cells, the durations of some cell cycle phases are underestimated.

### Cell Cycle Synchronization

150,000 cells were plated in 6 cm plates (for matched protein lysates and flow cytometry-based cell cycle analysis) or 6-well plates (for matched LCMS-based metabolite measurements and flow cytometry-based cell cycle analysis) in DMEM with 10% FBS. The following day, cells were washed three times with PBS and media was replaced with media containing RO-3306 (Selleckchem). 4.5-9 μM RO-3306 was used, with the concentration for different lots of RO-3306 adjusted to obtain optimal synchronization for experiments. After treating cells for 18 hours with RO-3306 (or DMSO for unsynchronized controls), cells were released from cell cycle arrest by washing three times with PBS and replacing the media with untreated media. Where relevant, media containing either 50 nM AZ20 (or DMSO as vehicle) was added at the time of release from RO-3306. Cells were collected at the indicated time points after release from arrest. For each experiment, parallel samples for each time point were analyzed by flow cytometry to assess cell cycle distribution.

### Data availability

The raw data associated with experiments presented in this study are available from the corresponding author upon request.

### Supplementary Video Files

**Supplementary Video 1. Example of untreated, cycling cells.**

Untreated A549 cells expressing the mVenus-Gem1 reporter. Imaging began at the start of the experiment and images were taken every 45 minutes thereafter.

**Supplementary Video 2. Example of cells that enter S phase after induction of nucleotide imbalance.**

Example of mVenus-Gem1-expressing A549 cells that were treated with 200 µM guanine and were in G1 phase at the time of guanine addition. Imaging began at the time of guanine addition and images were taken every 45 minutes thereafter.

**Supplementary Video 3. Example of cells that enter S phase before induction of nucleotide imbalance.**

Example of mVenus-Gem1-expressing A549 cells that were treated with 200 µM guanine and were in S/G2 phase at the time of guanine addition. Imaging began at the time of guanine addition and images were taken every 45 minutes thereafter.

