## Supplementary Data for "Nucleotide imbalance decouples cell growth from cell proliferation"

### Extended Data Figure 1

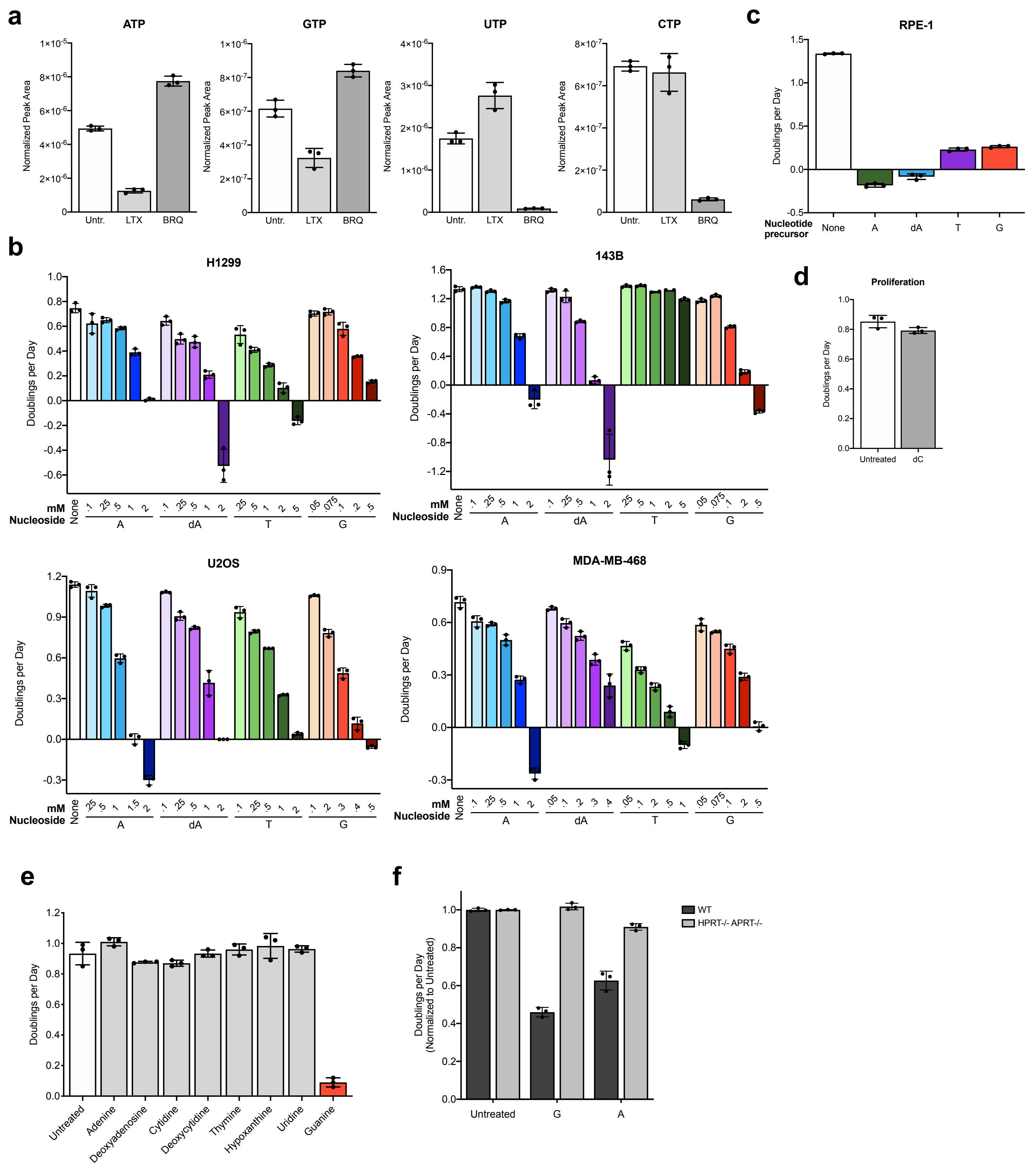

**Extended Data Figure 1. Salvage of different nucleobases and nucleosides can inhibit proliferation in different cell types.** **a**, Intracellular nucleotide levels in A549 cells cultured in standard conditions (Untr.), or treated with 1 μM lometrexol (LTX) or 1 μM brequinar (BRQ) as indicated. **b**, Proliferation rates of the indicated cell lines in standard culture conditions (None) or treated with the indicated concentrations of adenine (A), deoxyadenosine (dA), thymidine (T), or guanine (G). Of note, 143B cells are deficient for thymidine kinase, and therefore cannot salvage thymidine to produce dTMP. **c**, Proliferation rates of RPE-1 cells in standard culture (None) or treated with 1 mM T, 5 mM A, 1.5 mM dA, or 500 μM G. **d**, Proliferation rates of A549 cells in standard culture conditions (Untreated) or treated with 14 mM deoxycytidine (dC). **e**, Proliferation rates of A549 cells in standard culture conditions (Untreated) or treated with 200 μM of the indicated nucleobase/nucleoside. **f**, Normalized proliferation rates of A9 cells that are wild type (WT) or deficient (HPRT-/- APRT-/-) for hypoxanthine-guanine phosphoribosyltransferase and adenine phosphoribosyltransferase in standard culture conditions (Untreated) or treated with 200 μM G or A. Data are presented as mean +/- SD of 3 biological replicates.

### Extended Data Figure 2

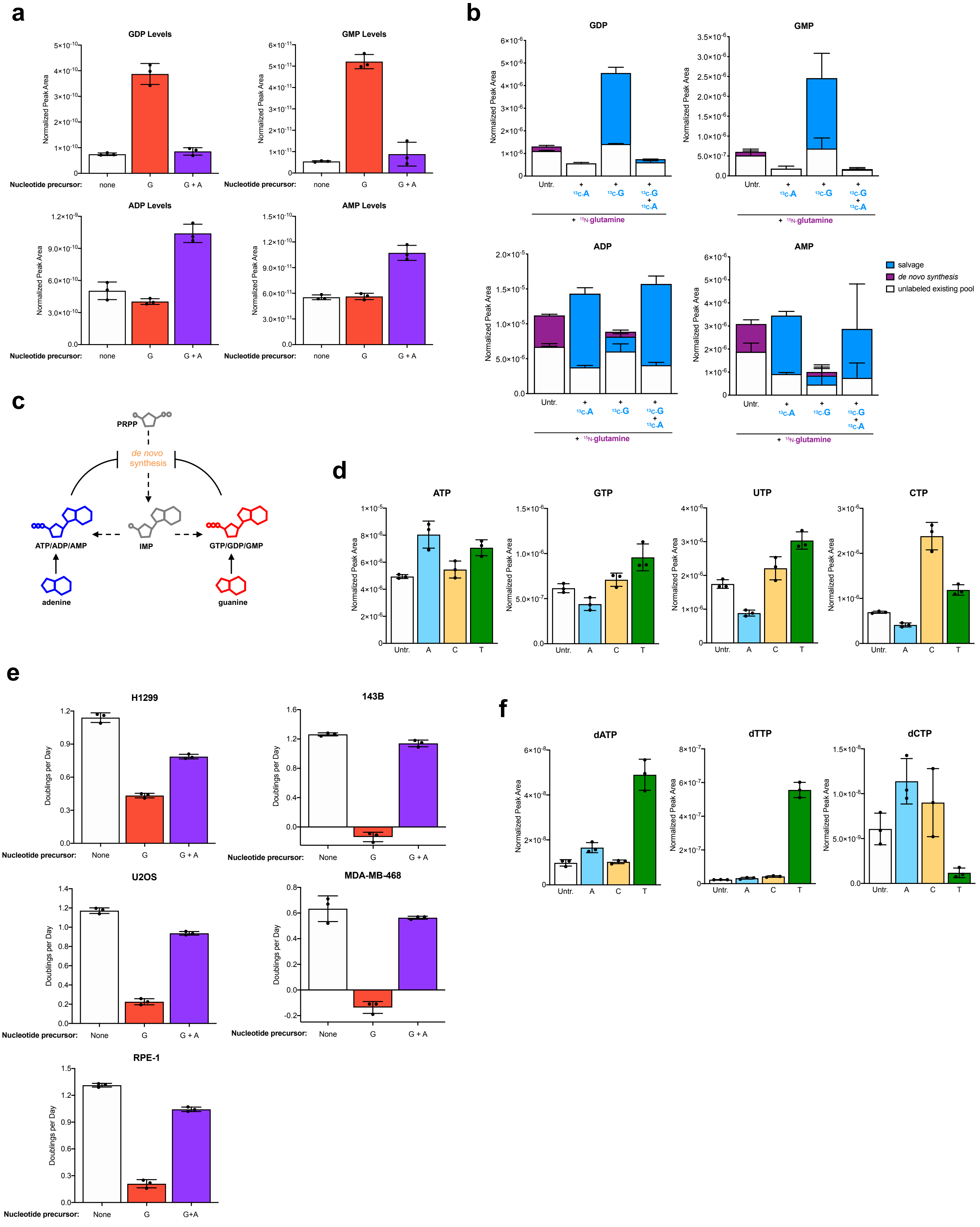

**Extended Data Figure 2. Salvage of excess nucleotides can induce nucleotide imbalance and impair proliferation.** **a**, Levels of the indicated nucleotides in A549 cells cultured in standard conditions (none) or treated for 24 hours with 200 μM guanine (G) with or without 200 μM adenine (A) as indicated. **b**, Total levels and labeling of the indicated nucleotides in A549 cells cultured for 24 hours in media containing $^{15}$N-amide-glutamine with or without 200 μM $^{13}$C-guanine ($^{13}$C-G) and/or $^{13}$C-adenine ($^{13}$C-A) as indicated. **c**, Diagram showing feedback regulation of purine synthesis. Adenylate and guanylate purines can allosterically inhibit enzymes involved in *de novo* purine synthesis. PRPP, phosphoribosyl pyrophosphate; IMP, inosine monophosphate. **d**, Intracellular nucleotide levels of A549 cells in standard culture conditions (Untr.) or treated with 200 μM A, 200 μM C, or 200 μM T as indicated. **e**, Proliferation rates of H1299, U2OS, MDA-MB-468, and RPE-1 cells in standard culture conditions (None) or treated with 200 μM G with or without 200 μM A, or 500 μM G with or without 500 μM A for RPE-1 cells, as indicated. **f**, Intracellular levels of dNTPs in A549 cells in standard culture conditions (Untreated) or treated with 200 μM A, 200 μM C, or 200 μM T as indicated. All nucleotide levels were measured using LCMS. Data are presented as mean +/- SD of 3 biological replicates.

### Extended Data Figure 3

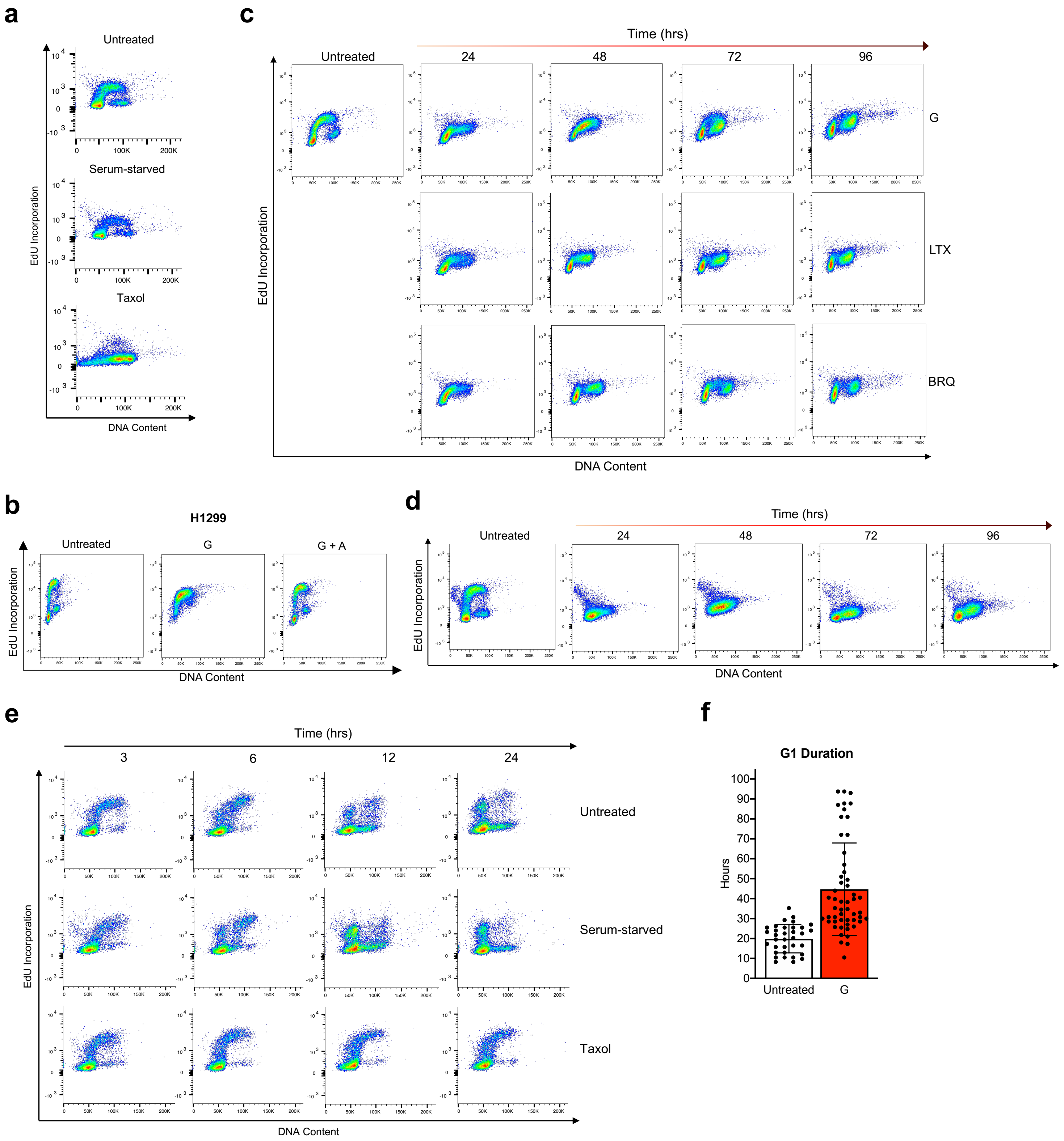

**Extended Data Figure 3. Nucleotide imbalance and depletion differentially alter cell cycle progression.** **a**, Cell cycle distribution as assessed by propidium iodide staining and EdU incorporation of A549 cells cultured for 24 hours in standard media (Untreated), in media lacking FBS (Serum-starved), or with 10 μM Taxol. Cells were pulsed with EdU for 30 minutes after each treatment and analyzed as outlined in Fig. 3a. **b**, Cell cycle distribution of H1299 cells cultured for 24 hours in standard conditions (Untreated) or treated with 500 μM guanine (G) with or without 500 μM adenine (A) as indicated. Cells were pulsed with EdU for 30 minutes after each treatment and analyzed as outlined in Fig. 3a. **c**, Cell cycle distribution of A549 cells cultured in standard conditions (Untreated) or treated for the indicated amount of time with 200 μM G, 1 μM lometrexol (LTX), or 1 μM brequinar (BRQ) as indicated. Cells were pulsed with EdU for 30 minutes after each treatment and analyzed as outlined in Fig. 3a. **d**, Cell cycle distribution of A549 cells cultured in standard conditions (Untreated) or treated with 1 mM thymidine for the indicated amount of time. As EdU is a thymidine analog, thymidine supplementation is expected to blunt EdU incorporation. Cells were pulsed with EdU for 30 minutes after each treatment and analyzed as outlined in Fig. 3a. **e**, Cell cycle distribution of A549 cells pulsed with EdU (see Fig. 3d) and then cultured for the indicated amount of time in standard conditions (Untreated), in media lacking FBS (Serum-starved), or with 10 μM Taxol as indicated. Note that the Untreated samples shown are from the same experiment shown in Fig. 3e. **f**, Duration of G1 phase in A549 cells expressing an mVenus-Gem1 reporter (see Fig. 3f) cultured in standard conditions (Untreated) or with 200 μM G, as assessed using live-cell imaging. 87 cells were analyzed. Data are presented as mean +/- SD.

### Extended Data Figure 4

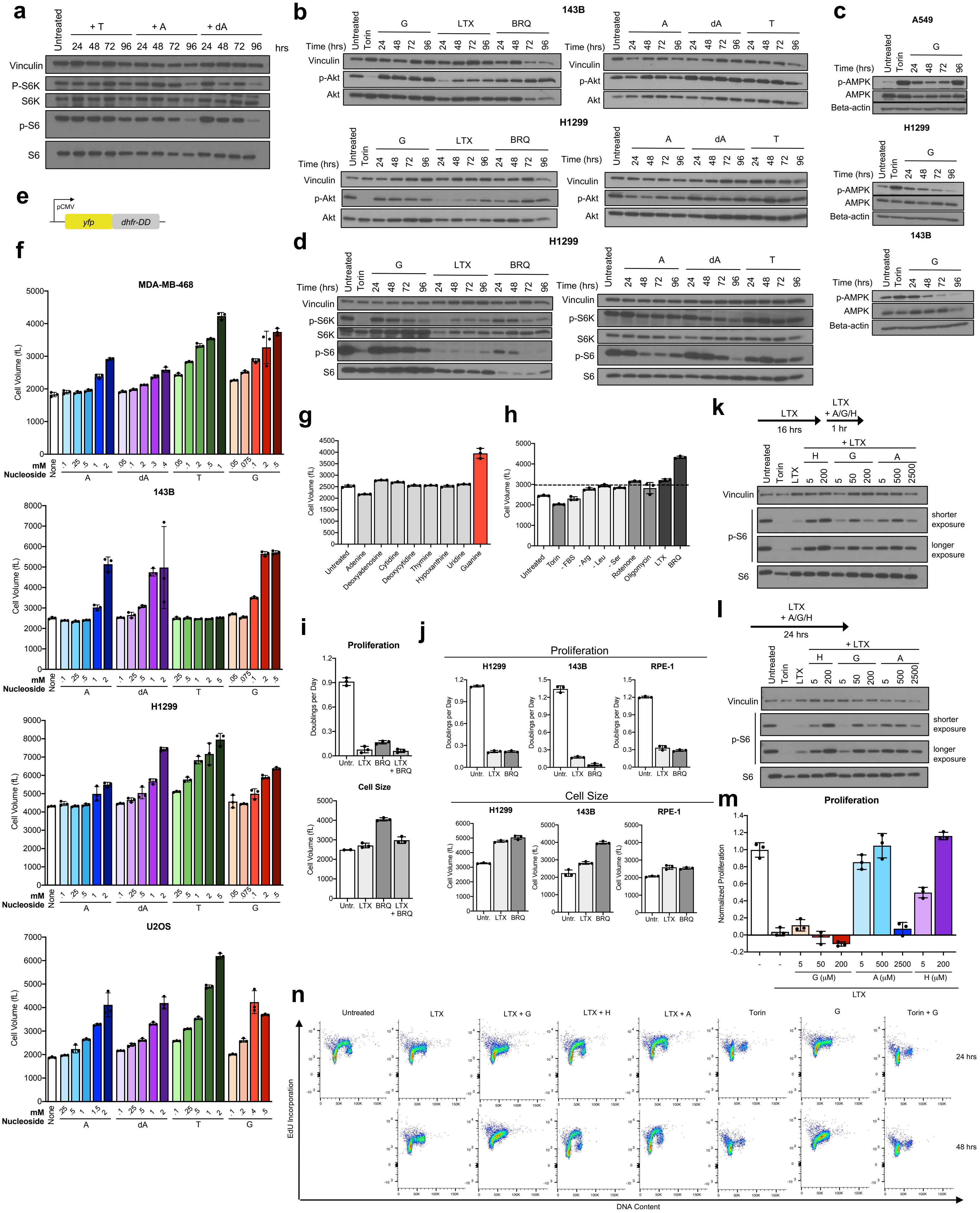

**Extended Data Figure 4. Under nucleotide imbalance, continued cell growth and S phase entry do not correlate with changes in growth signaling.** **a**, Western blot showing phosphorylation of ribosomal protein S6 and S6 kinase in A549 cells cultured in standard conditions (Untreated) or treated with 1mM thymidine (T), 2.5 mM adenine (A), or 1.5 mM deoxyadenosine (dA) for the indicated amount of time. Levels of vinculin, total S6K, and total S6 were also determined as controls. **b**, Western blots showing phosphorylation of Akt in 143B (top) and H1299 (bottom) cells cultured in standard conditions (Untreated) or treated with 1 µM Torin, 1 µM lometrexol (LTX), 1 µM brequinar (BRQ), 1 mM thymidine (T), 2.5 mM adenine (A), or 1.5 mM deoxyadenosine (dA) for the indicated amount of time. Levels of vinculin and total Akt were also determined as controls. **c**, Western blots showing phosphorylation of AMPK in A549, H1299, or 143B cells cultured in standard conditions (Untreated) or treated with 1 µM Torin or 200 µM G for the indicated amount of time. Levels of vinculin and total AMPK were also determined as controls. **d**, Western blots showing phosphorylation of ribosomal protein S6 and S6 kinase in H1299 cells cultured in standard conditions (Untreated) or treated with 1 µM Torin, 1 µM lometrexol (LTX), 1 µM brequinar (BRQ), 1 mM thymidine (T), 2.5 mM adenine (A), or 1.5 mM deoxyadenosine (dA) for the indicated amount of time. Levels of vinculin, total S6K, and total S6 were also determined as controls. **e**, Schematic showing design of a protein synthesis reporter where YFP is fused to an engineered unstable *E. coli* dihydrofolate reductase (DHFR) that acts as a degron. **f**, Mean volume of the indicated cell line cultured in standard conditions (None) or treated for 96 hours with the indicated concentration of A, dA, T, or guanine (G). Of note, 143B cells are deficient for thymidine kinase, and therefore cannot salvage thymidine to produce dTMP. **g**, Mean volume of A549 cells cultured in standard conditions (Untreated) or treated with 200 µM of the indicated nucleobase/nucleoside for 96 hours. **h**, Mean volume of A549 cells cultured for 96 hours in standard culture conditions (Untreated), or with 1 µM Torin1, without serum (-FBS), without arginine (-Arg), without leucine (-Leu), with 100 nM rotenone, with 5 nM oligomycin, with 1 µM LTX, or with 1 µM BRQ as indicated. **i**, Proliferation rate (top) and mean volume (bottom) of A549 cells cultured in standard conditions (Untr.) or treated for 96 hours with 1 µM lometrexol (LTX) or 1μM brequinar (BRQ) as indicated. **j**, Proliferation rate (left) and mean volume (right) of H1299, 143B, and RPE-1 cells cultured in standard conditions (Untr.) or treated for 96 hours with 1 μM lometrexol (LTX) or 1 μM brequinar (BRQ) as indicated. **k**, Phosphorylation of ribosomal protein S6 in A549 cells cultured for 16 hours in standard conditions (Untreated) or with 1 μM LTX, then supplemented for 1 hour with the indicated concentrations of G, A, or hypoxanthine (H). Levels of vinculin and total S6 were also determined as controls. **l**, Phosphorylation of ribosomal protein S6 in A549 cells cultured for 24 hours in standard conditions (Untreated) or treated with 1 μM LTX with or without the indicated concentrations of A, G, or H. Levels of vinculin and total S6 were also determined as controls. **m**, Proliferation rates of A549 cells cultured in media with or without 1 μM LTX, and with or without the indicated concentrations of G, A, or H. **n**, Cell cycle distribution of A549 cells cultured for 24 hours in standard conditions (Untreated), with 1 μM LTX with or without 200 μM G, 200 μM H, or 200 μM A as indicated, or with 10 μM Torin, 200 μM G, or both Torin and, as indicated. Cells were pulsed with EdU for 30 minutes after each treatment and then analyzed as outlined in Fig. 3a. Data are presented as mean +/- SD of 3 biological replicates.

#### Extended Data Figure 5

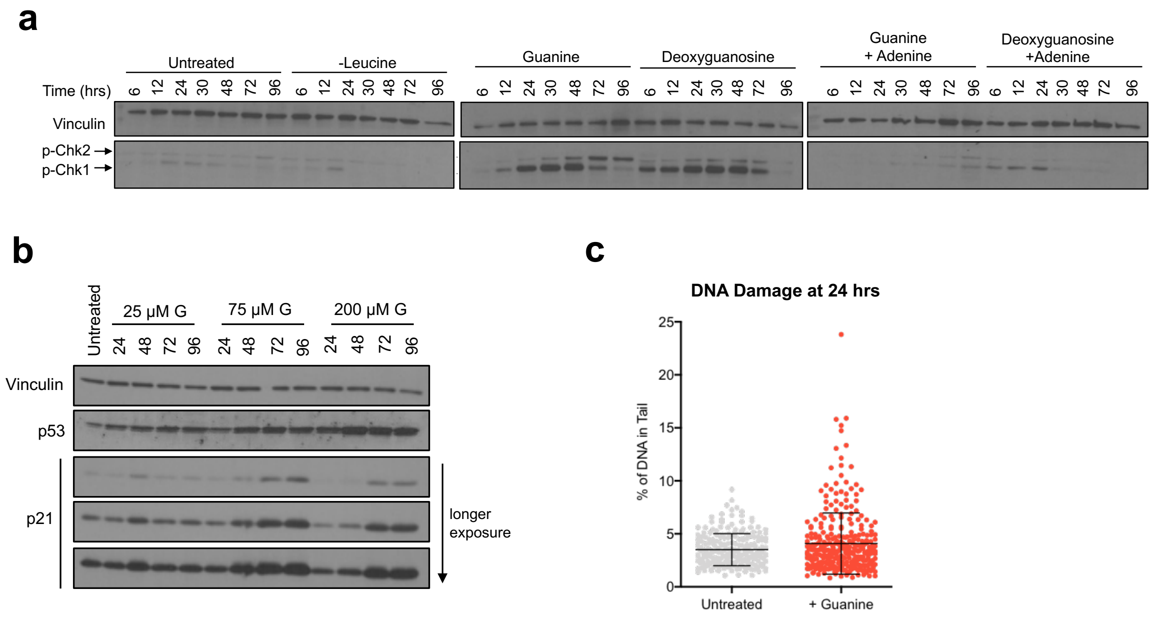


**Extended Data Figure 5. Imbalanced purines induce replication stress signaling. a**, Western blot showing phosphorylation of Chk1 and Chk2 in A549 cells cultured for the indicated time in standard media (Untreated) or media lacking leucine, or with addition of 200 μM guanine or 20 μM deoxyguanosine, with or without addition of 200 μM adenine as indicated. Levels of vinculin are also shown as a loading control. **b**, Western blot showing levels of p53 and p21 in A549 cells in standard media (Untreated) or cultured for the indicated amount of time with the indicated concentration of guanine (G). Levels of vinculin are also shown as a loading control. **c**, Comet assay to assess the presence of both single-stranded DNA and double-stranded DNA breaks in A549 cells treated without (Untreated) or with 200 μM guanine. 529 cells were analyzed. Data are presented as mean +/- SD.

### Extended Data Figure 6

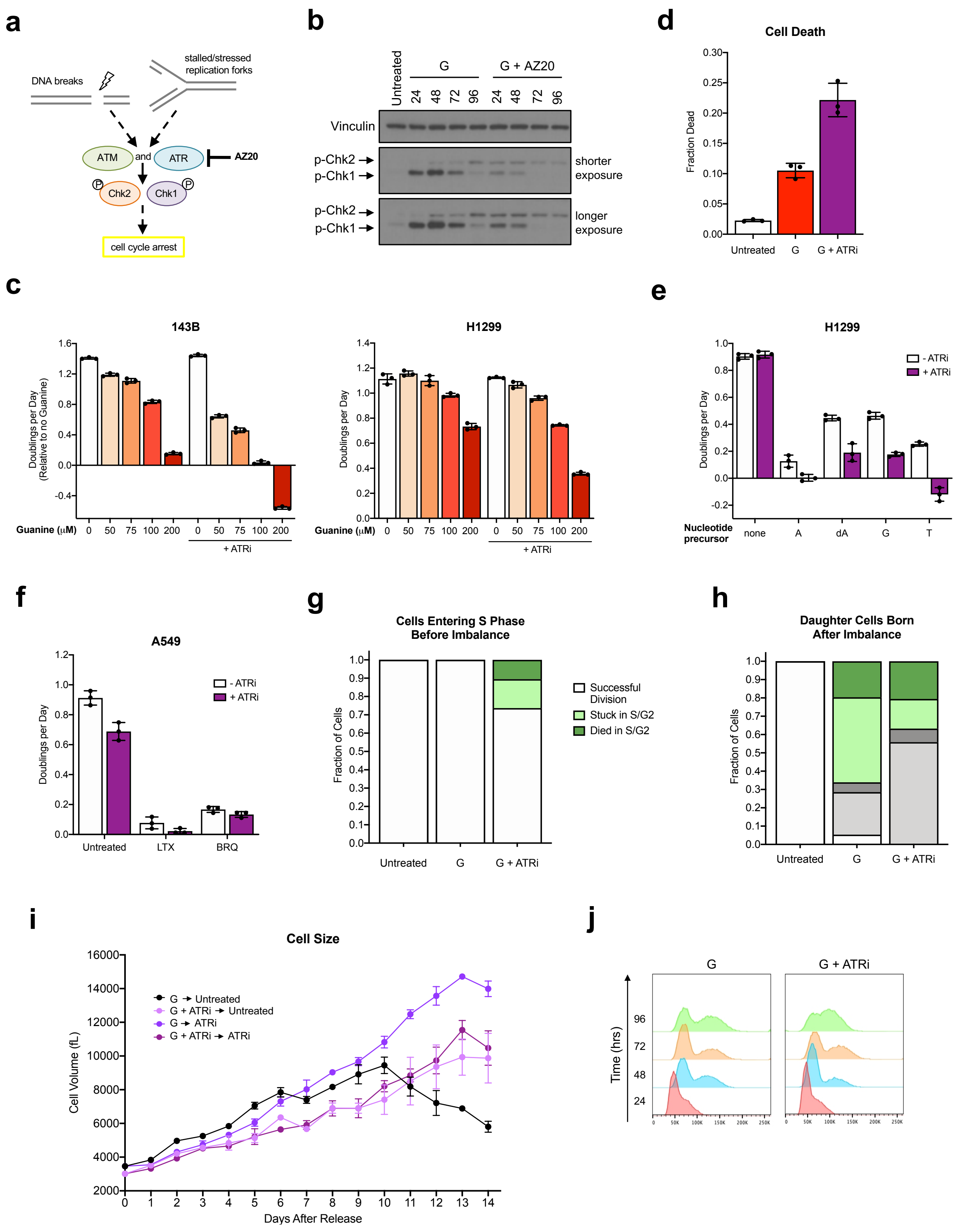

**Extended Data Figure 6. ATR signaling impacts fate of cells with imbalanced nucleotides. a**, Schematic outlining how ATR and ATM kinases respond to replication stress and DNA damage. The ATR and ATM targets Chk1 and Chk2 activate downstream effectors that halt cell cycle progression. AZ20 is an inhibitor of ATR kinase activity. **b**, Western blot showing phosphorylation of Chk1 and Chk2 in A549 cells cultured in standard media (Untreated) or treated for the indicated time with 200 μM guanine (G) with or without 50 nM AZ20. Levels of vinculin are also shown as a loading control. **c**, Proliferation rates of 143B and H1299 cells treated with the indicated concentration of guanine with or without 50 nM AZ20 (ATRi). **d**, Cell death measured in A549 cells cultured in standard conditions (Untreated) or treated with 200 μM G with or without 50 nM ATRi as indicated. **e**, Proliferation rate of H1299 cells cultured in standard conditions (none) or treated with 2 mM adenine (A), 1.5 mM deoxyadenosine (dA), 200 μM guanine (G), or 1 mM thymidine (T), with or without 50 nM ATRi as indicated. **f**, Proliferation rate of A549 cells cultured in standard conditions (Untreated) or treated with 1 μM lometrexol (LTX) or 1 μM brequinar (BRQ) with or without 50 nM ATRi as indicated. **g**, Cell fate as assessed using live-cell imaging of A549 mother cells expressing the mVenus-Gem1 reporter that were in S/G2 phase at the time of addition of 200 μM G with or without 50 nM ATRi. The fate of mother cells in S/G2 not exposed to excess G is also shown (Untreated). 83 cells were analyzed. **h**, Cell fate as assessed using live-cell imaging of A549 daughter cells expressing the mVenus-Gem1 reporter that were born after the addition of 200 μM G with or without 50 nM ATRi. The fate of daughter cells not exposed to excess G is also shown (Untreated). 158 cells were analyzed. **i**, Mean volume of A549 cells measured over time after release from treatment with G with or without ATRi as described in Fig. 6c. **j**, Cell cycle distribution as assessed by DNA content of A549 cells at 24-hour intervals after treatment with 200 μM G with or without 50 nM ATRi as indicated. Data are presented as mean +/- SD of 3 biological replicates.

### Extended Data Figure 7

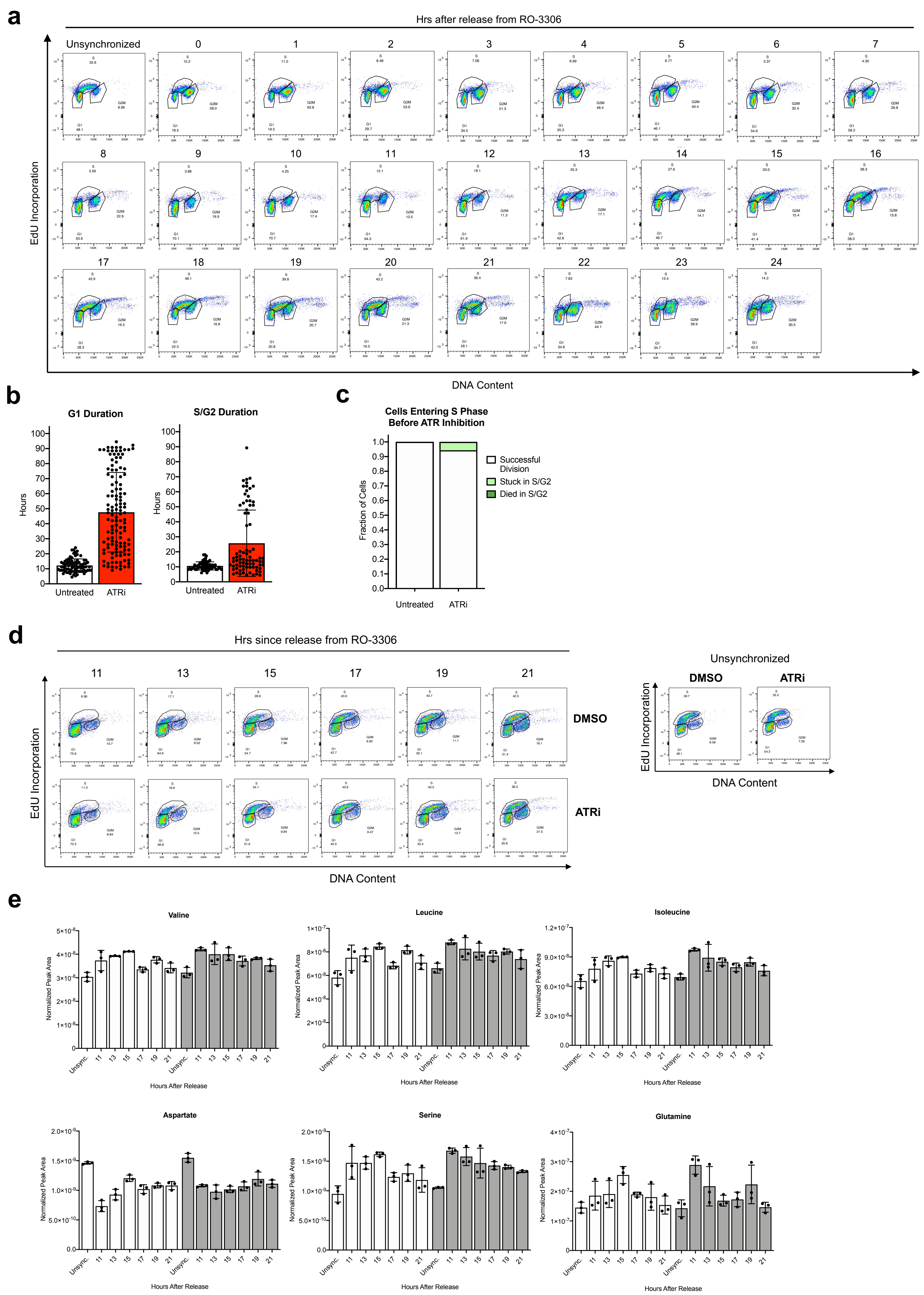

**Extended Data Figure 7. Amino acid levels remain relatively constant during unperturbed cell cycles, and are not impacted by loss of ATR signaling.** **a**, Cell cycle distribution of A549 cells corresponding to Western blots shown in Fig. 7b. Cells were arrested in G2 phase by treating with 9 µM RO-3306 for 18 hours, then RO-3306 was removed to release from cell cycle arrest for the indicated amount of time. Cells were pulsed with EdU for 30 minutes prior to each time point and analyzed as outlined in Fig. 3a. **b**, Duration of G1 phase and S/G2 phases in A549 cells expressing the mVenus-Gem1 reporter cultured in standard conditions (Untreated) or with 50 nM ATRi, as assessed using live-cell imaging. 87 cells were analyzed. Data are presented as mean +/- SD. **c**, Cell fate as assessed using live-cell imaging of A549 mother cells expressing the mVenus-Gem1 reporter that were in S/G2 phase when 50 nM AZ20 (ATRi) was added. The fate of mother cells in S/G2 not exposed to excess ATRi is also shown (Untreated). 55 cells were analyzed. **d**, Cell cycle distribution of A549 cells corresponding to the metabolite measurements shown in panel **e**, and in Fig. 7e and 7f. Cells were arrested in G2 phase by treating with 4.5 µM RO-3306 for 18 hours, then RO-3306 was removed to release from cell cycle arrest for the indicated amount of time. Cells were pulsed with EdU for 30 minutes prior to each time point and analyzed as outlined in Fig. 3a. Unsynchronized cells were treated with DMSO or 50 nM ATRi for 24 hours as indicated. **e**, Levels of the indicated amino acids in A549 cells synchronized in G2 phase by treating with 4.5 µM RO-3306 for 18 hours, then released into the cell cycle for the indicated amount of time. At the time of release from RO-3306, cells were either treated with DMSO or 50 nM ATRi as indicated. Unsynchronized cells (Unsync.) were treated with DMSO or 50 nM ATRi for 24 hours as indicated. All metabolite levels were measured by LCMS. Data are presented as mean +/- SD of 3 biological replicates.
